# Allosteric Cross-Talk Among SARS-CoV-2 Spike’s Receptor-Binding Domain Mutations Triggers an Effective Hijacking of Human Cell Receptor

**DOI:** 10.1101/2021.04.30.441093

**Authors:** Angelo Spinello, Andrea Saltalamacchia, Jure Borišek, Alessandra Magistrato

## Abstract

The rapid and relentless emergence of novel highly transmissible SARS-CoV-2 variants, possibly decreasing vaccine efficacy, currently represents a formidable medical and societal challenge. These variants frequently hold mutations on the Spike protein’s Receptor-Binding Domain (RBD), which, binding to the Angiotensin-Converting Enzyme 2 (ACE2) receptor, mediates viral entry into the host cells.

Here, all-atom Molecular Dynamics simulations and Dynamical Network Theory of the wild-type and mutant RBD/ACE2 adducts disclose that while the N501Y mutation (UK variant) enhances the Spike’s binding affinity towards ACE2, the N501Y, E484K and K417N mutations (South African variant) aptly adapt to increase SARS-CoV-2 propagation via a two-pronged strategy: (i) effectively grasping ACE2 through an allosteric signaling between pivotal RBD structural elements; and (ii) impairing the binding of antibodies elicited by infected/vaccinated patients. This information, unlocking the molecular terms and evolutionary strategies underlying the increased virulence of emerging SARS-CoV-2 variants, set the basis for developing the next-generation anti-COVID-19 therapeutics.

**TOC GRAPHICS:** 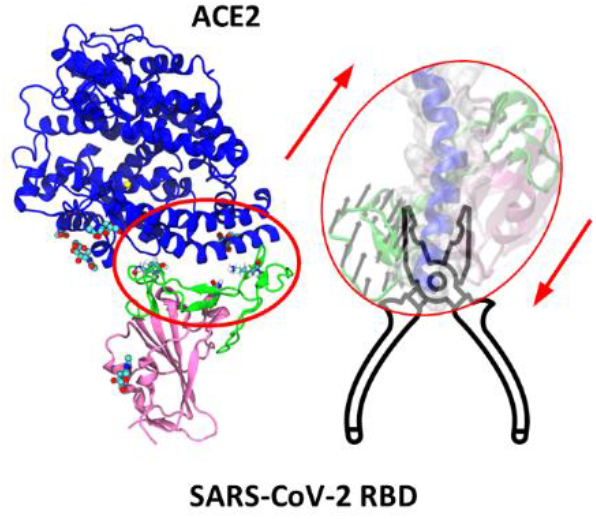

---

The severe acute respiratory syndrome coronavirus 2 (SARS-CoV-2), the etiological agent of coronavirus disease 19 (COVID-19), has infected as of April 30^rd^ about 150 million patients, causing over 3 million deaths worldwide. Owing to an unprecedentedly intense and relentless scientific effort, a variety of vaccines and monoclonal antibodies are becoming available for COVID-19 prophylaxis and therapeutic treatment.^1–3^

Alike other β-coronavirus (β-CoVs) the receptor-binding domain (RBD) of the homotrimeric viral spike (S) protein of SARS-CoV-2 mediates the molecular recognition and the binding to the human cellular receptor, angiotensin-converting enzyme 2 (ACE2),^4–5^ thus triggering SARS-CoV-2 entry into host cells. As such, the S-protein has been object of burgeoning research interest, becoming the prominent target for antibodies development. This prompted an exhaustive experimental^6–8^ and computational^9–12^ assessment of the molecular interactions between the S-protein and ACE2.

The worldwide continuous and uncontrolled transmission of SARS-CoV-2 set the condition for its rapid evolution into more infectious variants. As an example, one of the first S-protein mutation, D614G, characterized by an enhanced transmissibility, has rapidly become dominant.^13^ As well, other alarming strains have emerged: in United Kingdom (lineage B.1.1.7),^14^ South Africa (lineage B.1.351)^15^ and Brazil (lineage P.1),^16^ hereafter referred as the UK, SA and BR variants, respectively. Ultimately, a new dire Indian variant (lineage B.1.617) come to the fore. These lineages are object of raising concerns owing to their increased transmissibility and/or their potential ability to escape from infection/vaccines-induced immunity.

As concerns the most prominent nonsynonymous mutations placed in the S-protein’s RBD, most SARS-CoV-2 variants share the N501Y substitution (Figure 1), most likely implicated into an enhanced binding affinity towards ACE2,^17–19^ although preliminary reports indicates that this variant retains vaccine efficacy.^20^ In addition to N501Y, the SA variant also exhibits the E484K and K417N RBD mutations. E484, the most frequently mutated residue in COVID-19 patients, becomes E484K in the SA and BR and E484Q in the Indian strains. As well, mutation of K417, either to N or a T, is shared by the SA and BR variants, respectively. These mutations have been linked to viral escape from mAbs developed by vaccinated/infected patients.^2,21–22^

**Figure 1.**
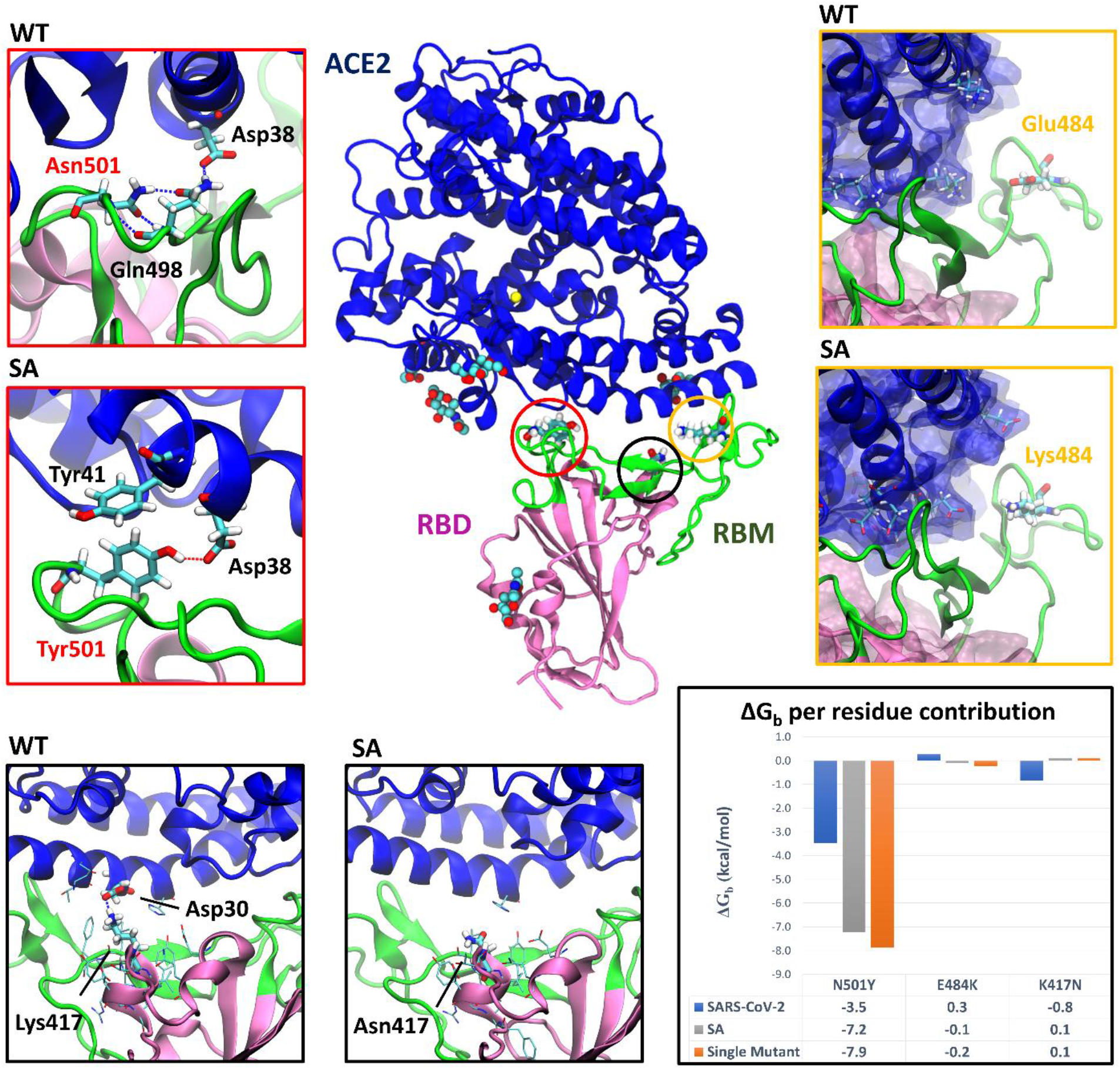
Representative structures of the complex between the South African (SA) SARS-CoV-2 variant of the receptor-binding domain (RBD, pink, with the receptor-binding motif (RBM) highlighted in green) and the angiotensin-converting enzyme 2 (ACE2, blue) as extracted from molecular dynamics simulations. The three N501Y, E484K and K417N mutations sites are circled in red, yellow, and black, respectively. The insets show a comparison of the key intermolecular interactions at the mutation sites in the wild-type and SA RBD/ACE2 complexes with residues depicted in licorice and hydrogen bonds displayed as dashed lines, along with the per-residue binding free energy (ΔG_b_) for the three mutations (bottom right).

Aiming to dissect at atomic-level the role of RDB mutations on the recognition of ACE2, we performed cumulative 15 μs all-atom molecular dynamics (MD) simulations of S-protein RBD/ACE2 complexes considering the RBD’s mutations present in the SA variant either concurrently or singularly.

Namely, we first built the adduct between ACE2 and RBD carrying the N501Y, E484K and K417N substitutions of the SA lineage (hereafter referred as ^SA^RBD/ACE2). Next, to inspect the role of each mutation, we built three distinct RBD/ACE2 models carrying N501Y (^N501Y^RBD/ACE2 or ^UK^RBD/ACE2), E484K (^E484K^RBD/ACE2) and K417N (^K417N^RBD/ACE2), ultimately comparing them with the WT RBD/ACE2 adduct (hereafter named RDB/ACE2). As a result, all the systems retain stable interactions at the RBD/ACE2 interface during 2.5 μs-long MD simulations (Figure S1). Most of the RBD residues binding to ACE2 lie within the receptor-binding motif (RBM), which is composed by two small β-strands and 4 flexible loops. In a previous study, we pinpointed the rigidity of RBD Loop3 (L3, composed by Thr470-Pro491) as the main factor underlying the larger binding affinity of SARS-CoV-2 towards ACE2,^9^ with respect to the closely-related SARS-CoV. In the current set of MD simulations all the investigated systems evidence an akin RBMs flexibility, with small differences being restricted to Loop1 and 4 (L1/4, Figure S2), where N501Y is placed (Figure 1).

Although not engaging direct interactions with ACE2, in the WT adduct N501@RBD intramolecularly H-bonds to Gln498, mediating the formation of a persistent H-bond network between the latter residue and Asp38@ACE2 (Figure 1), thus being the most dynamically correlated residue of the whole RBM (Figure S3).^9,23^ Nonetheless, in the ^SA^RBD/ACE2 and ^N501Y^RBD/ACE2 complexes N501Y further reinforces its prominent role in hijacking ACE2 by establishing π-stacking interactions with Tyr41@ACE2 (Figures 1, S4 and Table S1) and directly H-bonding to Asp38@ACE2, consistently with the ^N501Y^S-protein/ACE2 cryo-EM structure.^24^ Although the total binding free energies (ΔG_b_) of the distinct RBD/ACE2 adducts, calculated with the Molecular Mechanics/Generalized Born Surface Area (MM-GBSA) method,^25^ do not allow to discriminate the subtle differences between WT and mutant RDB/ACE2 adducts, the dissection of the per-residue amino acids ΔG_b_ contributions appears to be in agreement with experimental deep mutational scanning analyses.^18^ Indeed, the ΔG_b_ of Y501 increases with respect to its WT counterpart (−7.9 and −7.2 kcal/mol, in the ^N501Y^RBD/ACE2 and ^SA^RBD/ACE2 models, respectively) (Figure 1), while inducing a ΔG_b_ loss of nearby RBD residues (i.e. Glu498). The overall energetic gain of L1/4 region appears to be slightly favorable in ^N501Y^RBD/ACE2 and ^SA^RBD/ACE2, lowering the ΔG_b_ of 11-12 kcal/mol (Figure S4). As such, N501Y, present in the highly infective UK and SA variants, increases the RBD binding affinity for ACE2 (Figures 1, S4), consistently with experimental evidence.^18–19^ This comes along a more effective grasping and bending of the ACE2’s α1-helix in both ^N501Y^RBD/ACE2 and ^SA^RBD/ACE2 models as compared to RBD/ACE2 (Figure 2).

**Figure 2.**
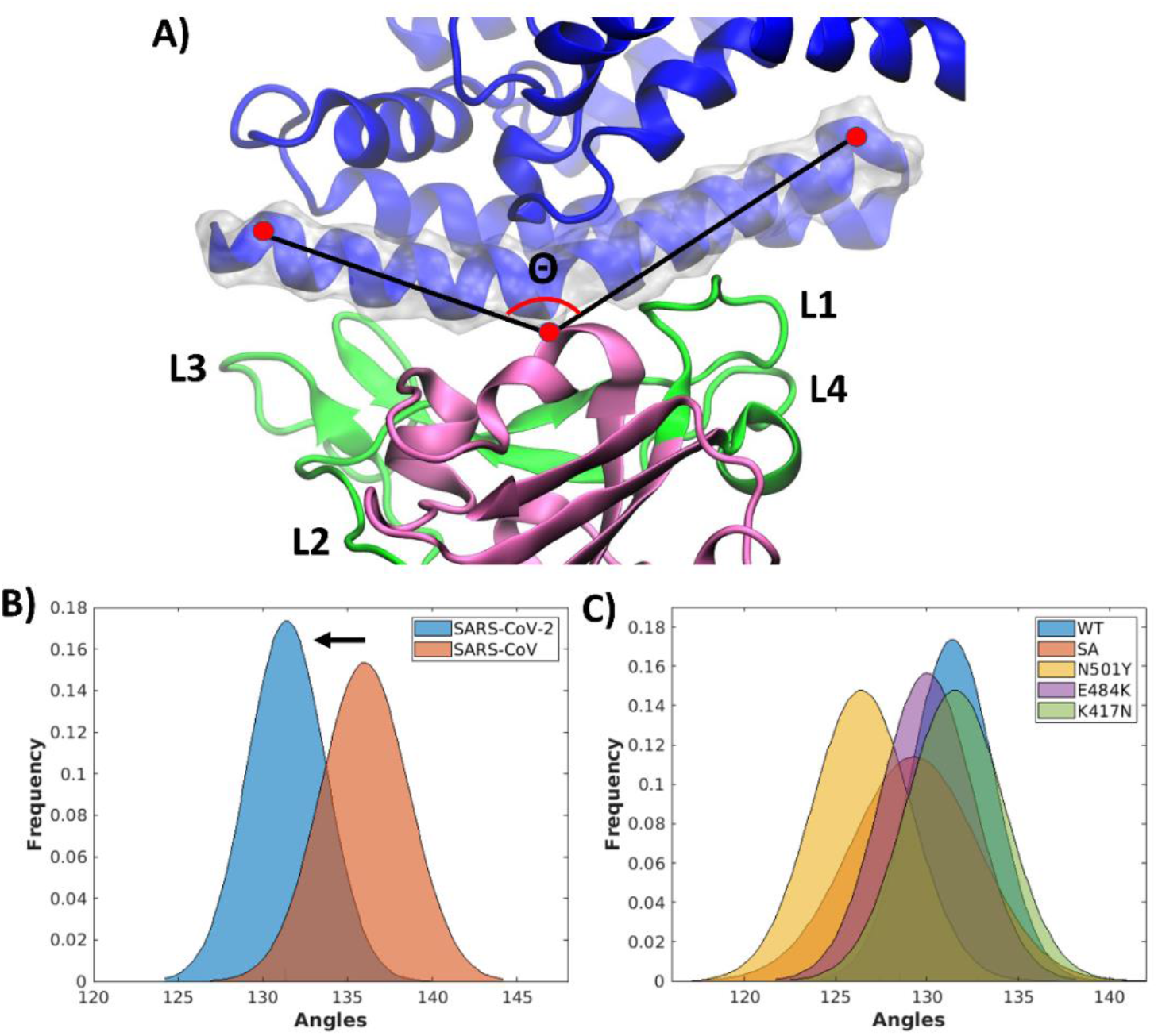
A) The bending angle (Θ) of the angiotensin-converting enzyme 2 (ACE2)’s α1-helix defined by the Cα atoms of Phe22, Asn53 and Trp69. The receptor-binding domain (RBD), motif (RBM) and ACE2 are displayed in pink, green and blue new-cartoons, respectively. The α1-helix@ACE2 is highlighted in silver transparent surface. B) Distribution of Θ angle (deg) for SARS-CoV-2 and SARS-CoV RBD/ACE2^9^ and C) for SARS-CoV-2 RBD/ACE2, ^SA^RDB/ACE2, ^N501Y^RDB/ACE2, ^E484K^RBD/ACE2 and ^K417N^RBD/ACE2 models.

We also inspected the role of E484K mutation common to the SA and BR variants. Consistently with deep mutational scanning,^18^ our MD simulations reveal that E484K only slightly increases the per-residue ΔG_b_ by 0.4±0.2 and 0.5±0.5 kcal/mol in ^SA^RBD/ACE2 and ^E484K^RBD/ACE2, respectively. Since E484K does not directly H-bonds to ACE2, the ΔG_b_ increase must owe to the electrostatic interactions between K484 and the nearby ACE2’s negatively charged (Asp and Glu) residues (Figure S5). Remarkably, K484 only modestly increases α1@ACE2 bending, as compared to the WT model (Figure 2C).

We finally assessed the role of K417,^15^ whose salt-bridge with Asp30@ACE2, present in half of the RBD/ACE2 MD trajectory, is lost upon K417N mutation (Figure S6 and Table S1). Consistently with experimental findings K417N triggers a small destabilization (ΔG_b_ decrease by 0.9±0.5 kcal/mol) of ^K417N^RBD/ACE2 and ^SA^RBD/ACE2 as compared to RBD/ACE2,^18^ and does not increase the α1-helix@ACE2 bending (Figure 2). Hence, the way K417N contributes to enhance the ACE2 sequestration remains elusive.

Due to the strategic location of K417, halfway of L1/4 and L3 in the RBM, tweezing the α1-helix@ACE2, we computed the cross-correlation matrices based on the Pearson’s correlation coefficient and the per-residue sum of the Cross-Correlation coefficient (CCc) for the residues at the RBM/ACE2 interface (Figure S3).^26–27^ Stunningly, ^SA^RBD/ACE2 and ^K417^RBD/ACE2 exhibit the largest CCcs all over the RBM, with the difference of ^SA^RBD/ACE2 being more marked at the L1/4 and L3 regions.

Aiming to assess whether the mutations could interfere with the RBD’s slow motions, we performed principal component (PC) analysis of WT and mutant RBD models to gather their most relevant motions (essential dynamics). As a result, PC1 and 2 of WT or all mutant RBD systems reveal an opening/closing motions of L1/4 and L3 regions, implicated in grasping α1-helix@ACE2 (Figure S7). Since an allosteric communication among SARS-CoV-2 mutations has been recently speculated,^28–30^ we then applied dynamical NetWork theory Analysis (NWA) to decrypt the information-exchange pathways underlying the observed RBD functional dynamics and to decode whether RBD mutants can enhance the allosteric cross-talk between critical RDB’s structural elements.^26,31^ In NWA, the protein is represented as a correlation-based weighted network. The nodes (the residues’ center of mass) are connected by edges whose numerical value (weight) indicates the correlation-strength between residues pairs (i.e. small/large weights reflect highly/poorly correlated and anticorrelated motions). By computing cross-correlations between residues along an MD trajectory, NWA finds the optimal and suboptimal signaling-paths between two user-selected source (484@L3) and sink (501@L4) residues. The outcoming path-lengths are thus inversely proportional to the signaling-strength and to the amount of correlation existing among their tracing nodes.^31^

By performing NWA on the RBD alone we observed several cross-communication paths crossing the RBM, which in RBD/ACE2 minorly involve even K417 (Figure 3B). Remarkably, in ^N501Y(UK)^RBD and ^SA^RBD these paths are shorter (the residues are more correlated) suggesting a stronger signaling between the two RBM extremities, which may result in a more effective opening/closing of the L1/4 and L3.

**Figure 3.**
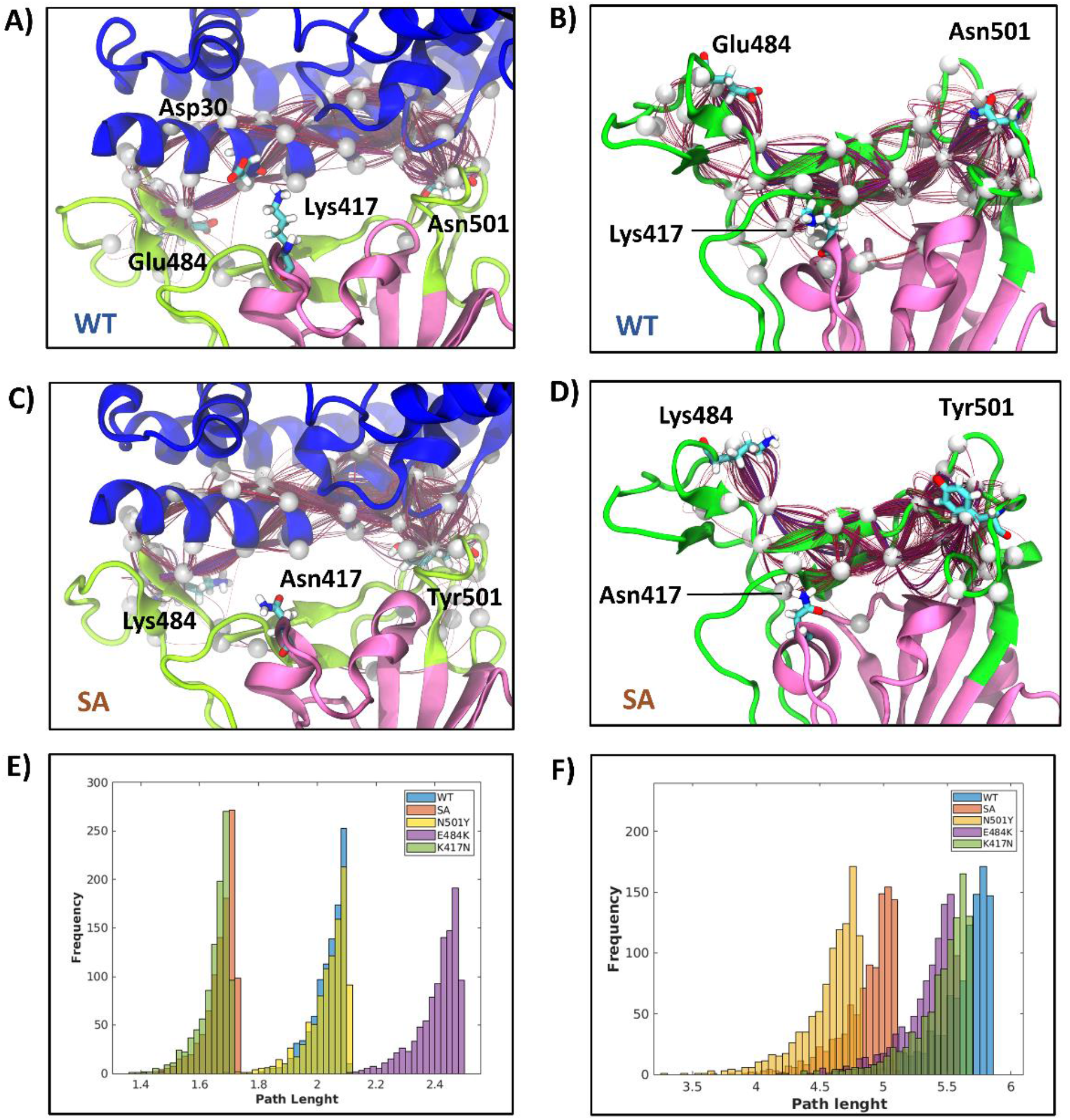
Optimal and suboptimal signaling-paths (red lines, with nodes depicted as white spheres) connecting the receptor-binding domain (RBD) residues 484 and 501, for A) wild type (WT) RBD/angiotensin-converting enzyme 2 (ACE2), B) WT RBD, C) South African ^SA^RBD/ACE2 and D) ^SA^RBD. Distribution of signaling-path lengths for E) all investigated RBD/ACE2 adducts and F) the isolated RBDs.

Stunningly, in all RBD/ACE2 models allosteric signaling within RBM occurs along the α1-helix@ACE2 (Figures 3A,C) and the path-lengths distribution of both ^SA^RBD/ACE2 and ^K417N^RBD/ACE2 (Figure 3E) is shifted towards lower values, suggesting that the motions of RBM’s residues are more tightly correlated, thus triggering a more effective ACE2 hijacking. The similar distribution observed for ^SA^RBD/ACE2 and ^K417N^RBD/ACE2 suggests that K417N is primarily liable for the enhanced cross-talk between critical RBD recognition loops. Conversely, the N501Y and E484K mutations does not affect and even contrast this cross-talk, respectively (Figure 3E).

To identify the residues critically involved in the signaling-pathways we computed the node degeneracy (i.e. number of times a node is present in the calculated paths). In the presence of the RBD mutations a significant variation in degeneracy is observed for those residues engaging H-bond or hydrophobic interactions at the RBD/ACE2 interface, among which Asp38@ACE2, Asp355@ACE2, and Thr500@RBD (Figure S8 and Table S3). To further dissect the source of the increased cross-talk in the mutant RBD/ACE2 complexes we inspected whether RBD mutations alter the intra-RBD H-bonds networks at RBD/ACE2 interface. Interestingly, the main differences among the investigated systems are localized on L3-4, nearby the mutation sites (Table S4). In particular, a decrease of the intramolecular H-bond persistence of Asn487, which strongly H-bonds ACE2 (Table S1), results into a higher node-degeneracy in the signaling-route (Table S3).

Complementarily, structural X-ray and Cryo-EM studies elucidated that K417 and E484 RBD residues establish H-bonds with distinct mAbs isolated from COVID-19 patients’ sera. Hence, K417N and E484K substitutions alter the electrostatic complementarity between the RBD and class 1 and 2 mAbs, respectively (Figures 4 and Table S5),^32–37^ impairing mAbs binding and contributing to viral escape from vaccine/disease-induced immunity.^2,33^

**Figure 4.**
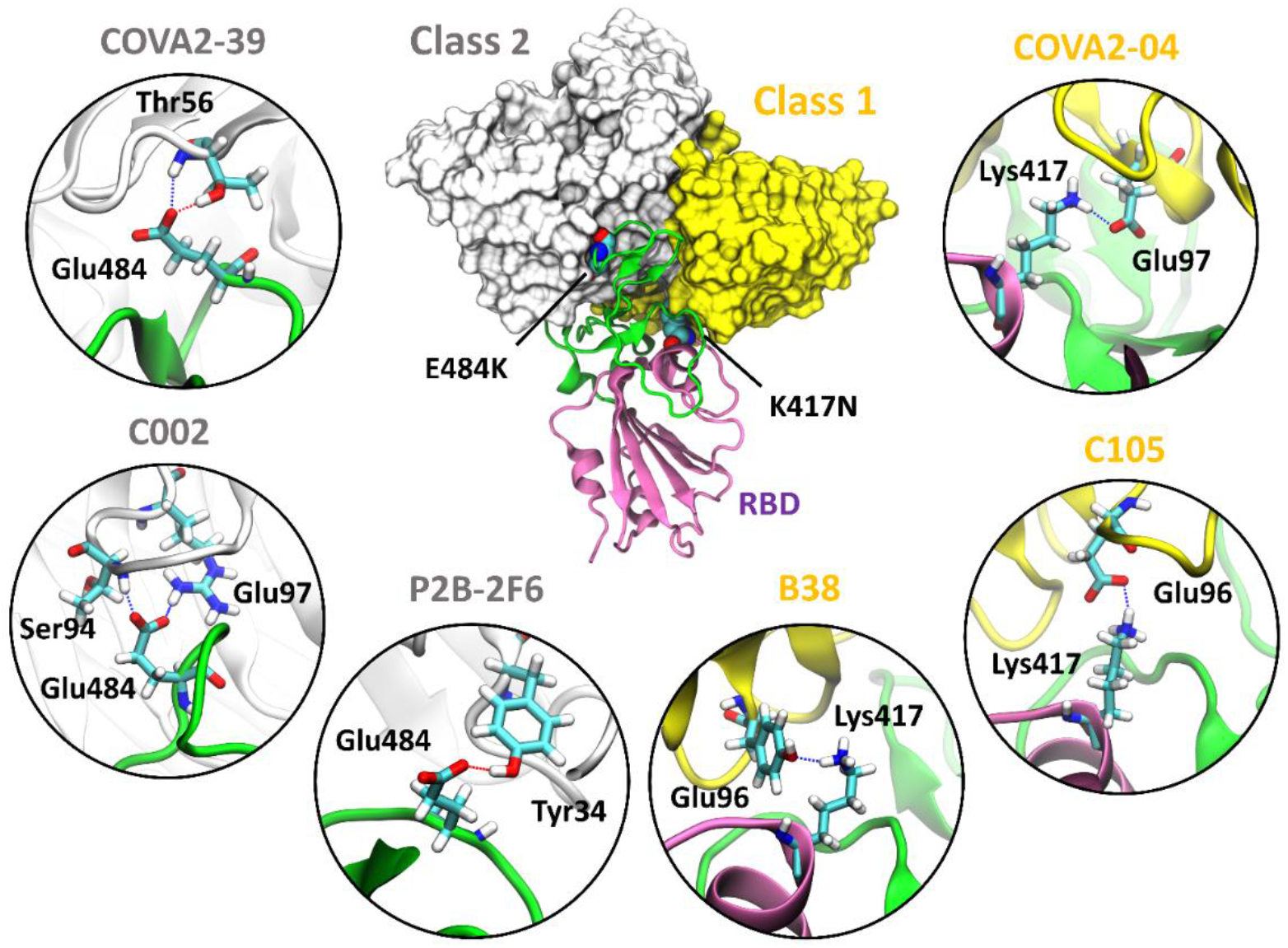
Binding mode of class 1 and class 2 (gray and yellow surfaces) monoclonal antibodies (mAbs) isolated from patients to the Spike’s receptor-binding domain (RBD), showing the use of different epitopes. The receptor-binding motif (RBM) and RBD are shown as green and pink new-cartoons, respectively. Insets disclose the key intermolecular interactions established between (from left to right) the mAbs COVA2-39,^32^ C002,^33^ P2B-2F6,^34^ B38,^35^ C105,^36^ and COVA2-04.^32^

In summary, aiming to dissecting the molecular basis for the higher infectivity and transmissibility of emerging SARS-CoV-2 variants, we have assessed the impact of the SA set of mutations, localized in the RBD, even inspecting the role of each single mutant. As a result, we disclose that while N501Y (hallmark of the UK variant) enhances the binding affinity towards ACE2 and α1-helix@ACE2 bending, the SA strain exploits a two-pronged strategy to more effectively infect the host cells by (i) increasing the allosteric signaling among the pivotal RBM extremes, which acting as a tweezer, more effectively grasp/bend the α1-helix@ACE2, (ii) destabilizing/impairing the interactions with class 1 and 2 mAbs (K417N and E484K, respectively) extracted from COVID-19 patients’ sera (Figure 4 and Table S5). Stunningly, the main actor in modulating the allosteric cross-talk among the RBD mutants appears to be K417N, whose role has remained so far elusive. In this scenario, it is tempting to argue that the BR variant, differing from the SA one only by the K417T@RBD substitution, may exploit the same strategy to foster viral propagation. Our outcomes contribute to decrypt at atomic-level the evolutionary strategies underlying the increased SARS-CoV-2 infectivity and spreading of emerging variants, setting a conceptual basis to devise the next-generation therapeutic strategies against current and future viral strains.

## ASSOCIATED CONTENT

Computational Details, Figure S1 to S8 and Table S1 to S5. This material is available free of charge at via the Internet at http://pubs.acs.org.

## ACKNOWLEDGMENT

AS was supported by a FIRC-AIRC “Mario e Valeria Rindi” fellowship for Italy. AM thanks the financial support of the Italian Association for Cancer Research (AIRC) (IG grant 24514) and of the project ‘Against bRain cancEr: finding personalized therapies with in Silico and in vitro strategies’ (ARES) CUP:D93D19000020007 POR FESR 2014 2020-1.3.b-Friuli Venezia Giulia.

## REFERENCES

(1) Hsieh, C. L.; Goldsmith, J. A.; Schaub, J. M.; DiVenere, A. M.; Kuo, H. C.; Javanmardi, K.; Le, K. C.; Wrapp, D.; Lee, A. G.; Liu, Y.; Chou, C. W.; Byrne, P. O.; Hjorth, C. K.; Johnson, N. V.; Ludes-Meyers, J.; Nguyen, A. W.; Park, J.; Wang, N.; Amengor, D.; Lavinder, J. J.; Ippolito, G. C.; Maynard, J. A.; Finkelstein, I. J.; McLellan, J. S. Structure-based design of prefusion-stabilized SARS-CoV-2 spikes. Science 2020, 369 (6510), 1501–1505.

(2) Andreano, E.; Piccini, G.; Licastro, D.; Casalino, L.; Johnson, N. V.; Paciello, I.; Monego, S. D.; Pantano, E.; Manganaro, N.; Manenti, A.; Manna, R.; Casa, E.; Hyseni, I.; Benincasa, L.; Montomoli, E.; Amaro, R. E.; McLellan, J. S.; Rappuoli, R. SARS-CoV-2 escape in vitro from a highly neutralizing COVID-19 convalescent plasma. bioRxiv 2020.

(3) Serapian, S. A.; Marchetti, F.; Triveri, A.; Morra, G.; Meli, M.; Moroni, E.; Sautto, G. A.; Rasola, A.; Colombo, G. The Answer Lies in the Energy: How Simple Atomistic Molecular Dynamics Simulations May Hold the Key to Epitope Prediction on the Fully Glycosylated SARS-CoV-2 Spike Protein. J. Phys. Chem. Lett. 2020, 11 (19), 8084–8093.

(4) Li, W.; Moore, M. J.; Vasilieva, N.; Sui, J.; Wong, S. K.; Berne, M. A.; Somasundaran, M.; Sullivan, J. L.; Luzuriaga, K.; Greenough, T. C.; Choe, H.; Farzan, M. Angiotensin-converting enzyme 2 is a functional receptor for the SARS coronavirus. Nature 2003, 426 (6965), 450–4.

(5) Wang, Q.; Zhang, Y.; Wu, L.; Niu, S.; Song, C.; Zhang, Z.; Lu, G.; Qiao, C.; Hu, Y.; Yuen, K. Y.; Wang, Q.; Zhou, H.; Yan, J.; Qi, J. Structural and Functional Basis of SARS-CoV-2 Entry by Using Human ACE2. Cell 2020, 181 (4), 894–904 e9.

(6) Wrapp, D.; Wang, N.; Corbett, K. S.; Goldsmith, J. A.; Hsieh, C. L.; Abiona, O.; Graham, B. S.; McLellan, J. S. Cryo-EM structure of the 2019-nCoV spike in the prefusion conformation. Science 2020, 367 (6483), 1260–1263.

(7) Shang, J.; Ye, G.; Shi, K.; Wan, Y.; Luo, C.; Aihara, H.; Geng, Q.; Auerbach, A.; Li, F. Structural basis of receptor recognition by SARS-CoV-2. Nature 2020.

(8) Tai, W.; He, L.; Zhang, X.; Pu, J.; Voronin, D.; Jiang, S.; Zhou, Y.; Du, L. Characterization of the receptor-binding domain (RBD) of 2019 novel coronavirus: implication for development of RBD protein as a viral attachment inhibitor and vaccine. Cell Mol. Immunol. 2020.

(9) Spinello, A.; Saltalamacchia, A.; Magistrato, A. Is the Rigidity of SARS-CoV-2 Spike Receptor-Binding Motif the Hallmark for Its Enhanced Infectivity? Insights from All-Atom Simulations. J. Phys. Chem. Lett. 2020, 11 (12), 4785–4790.

(10) Wang, Y.; Liu, M.; Gao, J. Enhanced receptor binding of SARS-CoV-2 through networks of hydrogen-bonding and hydrophobic interactions. Proc Natl Acad Sci U S A 2020, 117 (25), 13967–13974.

(11) Ali, A.; Vijayan, R. Dynamics of the ACE2-SARS-CoV-2/SARS-CoV spike protein interface reveal unique mechanisms. Sci Rep 2020, 10 (1), 14214.

(12) Frances-Monerris, A.; Hognon, C.; Miclot, T.; Garcia-Iriepa, C.; Iriepa, I.; Terenzi, A.; Grandemange, S.; Barone, G.; Marazzi, M.; Monari, A. Molecular Basis of SARS-CoV-2 Infection and Rational Design of Potential Antiviral Agents: Modeling and Simulation Approaches. J. Proteome Res. 2020, 19 (11), 4291–4315.

(13) Korber, B.; Fischer, W. M.; Gnanakaran, S.; Yoon, H.; Theiler, J.; Abfalterer, W.; Hengartner, N.; Giorgi, E. E.; Bhattacharya, T.; Foley, B.; Hastie, K. M.; Parker, M. D.; Partridge, D. G.; Evans, C. M.; Freeman, T. M.; de Silva, T. I.; Sheffield, C.-G. G.; McDanal, C.; Perez, L. G.; Tang, H.; Moon-Walker, A.; Whelan, S. P.; LaBranche, C. C.; Saphire, E. O.; Montefiori, D. C. Tracking Changes in SARS-CoV-2 Spike: Evidence that D614G Increases Infectivity of the COVID-19 Virus. Cell 2020, 182 (4), 812–827 e19.

(14) Davies, N. G.; Abbott, S.; Barnard, R. C.; Jarvis, C. I.; Kucharski, A. J.; Munday, J. D.; Pearson, C. A. B.; Russell, T. W.; Tully, D. C.; Washburne, A. D.; Wenseleers, T.; Gimma, A.; Waites, W.; Wong, K. L. M.; van Zandvoort, K.; Silverman, J. D.; Group, C. C.-W.; Consortium, C.-G. U.; Diaz-Ordaz, K.; Keogh, R.; Eggo, R. M.; Funk, S.; Jit, M.; Atkins, K. E.; Edmunds, W. J. Estimated transmissibility and impact of SARS-CoV-2 lineage B.1.1.7 in England. Science 2021, 372 (6538).

(15) Tegally, H.; Wilkinson, E.; Giovanetti, M.; Iranzadeh, A.; Fonseca, V.; Giandhari, J.; Doolabh, D.; Pillay, S.; San, W. J.; Msoni, N.; Mlisana, K.; von Gottberg, A.; Walaza, S.; Allam, M.; Ismail, A.; Mohale, T.; Glass, A. J.; Engelbrecht, S.; Van Zyl, G.; Preiser, W.; Petruccione, F.; Sigal, A.; Hardie, D.; Marais, G.; Hsiao, M.; Korsman, S.; Davies, M.; Tyers, L.; Mudau, I.; York, D.; Maslo, C.; Goedhals, D.; Abrahams, S.; Laguda-Akingba, O.; Alisoltani-Dehkordi, A.; Godzik, A.; Wibmer, C. K.; Sewell, B. T.; Lourenco, J.; Alcantara, L. C.; Pond, S. L. K.; Weaver, S.; Martin, D.; Lessells, R. J.; Bhiman, J. N.; Williamson, C.; de Oliveira, T. Emergence and rapid spread of a new severe acute respiratory syndrome-related coronavirus 2 (SARS-CoV-2) lineage with multiple spike mutations in South Africa. medRxiv 2020.

(16) He, D.; Fan, G.; Wang, X.; Li, Y.; Peng, Z. Emergence and rapid spread of a new severe acute respiratory syndrome-related coronavirus 2 (SARS-CoV-2) lineage with multiple spike mutations in South Africa. medRxiv 2021.

(17) Gu, H.; Chen, Q.; Yang, G.; He, L.; Fan, H.; Deng, Y. Q.; Wang, Y.; Teng, Y.; Zhao, Z.; Cui, Y.; Li, Y.; Li, X. F.; Li, J.; Zhang, N. N.; Yang, X.; Chen, S.; Guo, Y.; Zhao, G.; Wang, X.; Luo, D. Y.; Wang, H.; Yang, X.; Li, Y.; Han, G.; He, Y.; Zhou, X.; Geng, S.; Sheng, X.; Jiang, S.; Sun, S.; Qin, C. F.; Zhou, Y. Adaptation of SARS-CoV-2 in BALB/c mice for testing vaccine efficacy. Science 2020, 369 (6511), 1603–1607.

(18) Starr, T. N.; Greaney, A. J.; Hilton, S. K.; Ellis, D.; Crawford, K. H. D.; Dingens, A. S.; Navarro, M. J.; Bowen, J. E.; Tortorici, M. A.; Walls, A. C.; King, N. P.; Veesler, D.; Bloom, J. D. Deep Mutational Scanning of SARS-CoV-2 Receptor Binding Domain Reveals Constraints on Folding and ACE2 Binding. Cell 2020, 182 (5), 1295–1310 e20.

(19) Tian, F.; Tong, B.; Sun, L.; Shi, S.; Zheng, B.; Wang, Z.; Dong, X.; Zheng, P. Mutation N501Y in RBD of Spike Protein Strengthens the Interaction between COVID-19 and its Receptor ACE2. BioRxiv 2021.

(20) Xie, X.; Liu, Y.; Liu, J.; Zhang, X.; Zou, J.; Fontes-Garfias, C. R.; Xia, H.; Swanson, K. A.; Cutler, M.; Cooper, D.; Menachery, V. D.; Weaver, S. C.; Dormitzer, P. R.; Shi, P. Y. Neutralization of SARS-CoV-2 spike 69/70 deletion, E484K and N501Y variants by BNT162b2 vaccine-elicited sera. Nat. Med. 2021.

(21) Greaney, A. J.; Starr, T. N.; Gilchuk, P.; Zost, S. J.; Binshtein, E.; Loes, A. N.; Hilton, S. K.; Huddleston, J.; Eguia, R.; Crawford, K. H. D.; Dingens, A. S.; Nargi, R. S.; Sutton, R. E.; Suryadevara, N.; Rothlauf, P. W.; Liu, Z.; Whelan, S. P. J.; Carnahan, R. H.; Crowe, J. E., Jr.; Bloom, J. D. Complete Mapping of Mutations to the SARS-CoV-2 Spike Receptor-Binding Domain that Escape Antibody Recognition. Cell Host Microbe 2021, 29 (1), 44–57 e9.

(22) Weisblum, Y.; Schmidt, F.; Zhang, F.; DaSilva, J.; Poston, D.; Lorenzi, J. C.; Muecksch, F.; Rutkowska, M.; Hoffmann, H. H.; Michailidis, E.; Gaebler, C.; Agudelo, M.; Cho, A.; Wang, Z.; Gazumyan, A.; Cipolla, M.; Luchsinger, L.; Hillyer, C. D.; Caskey, M.; Robbiani, D. F.; Rice, C. M.; Nussenzweig, M. C.; Hatziioannou, T.; Bieniasz, P. D. Escape from neutralizing antibodies by SARS-CoV-2 spike protein variants. Elife 2020, 9.

(23) Pavlin, M.; Spinello, A.; Pennati, M.; Zaffaroni, N.; Gobbi, S.; Bisi, A.; Colombo, G.; Magistrato, A. A Computational Assay of Estrogen Receptor alpha Antagonists Reveals the Key Common Structural Traits of Drugs Effectively Fighting Refractory Breast Cancers. Sci. Rep. 2018, 8 (1), 649.

(24) Zhu, X.; Mannar, D.; Srivastava, S. S.; Berezuk, A. M.; Demers, J.; Saville, J. W.; Leopold, K.; Li, W.; Dimitrov, D. S.; Tuttle, K. S.; Zhou, S.; Chittori, S.; Subramaniam, S. Cryo-EM Structures of the N501Y SARS-CoV-2 Spike Protein in Complex with ACE2 and Two Potent Neutralizing Antibodies. BioRxiv 2021.

(25) Kollman, P. A.; Massova, I.; Reyes, C.; Kuhn, B.; Huo, S.; Chong, L.; Lee, M.; Lee, T.; Duan, Y.; Wang, W.; Donini, O.; Cieplak, P.; Srinivasan, J.; Case, D. A.; Cheatham, T. E., 3rd. Calculating structures and free energies of complex molecules: combining molecular mechanics and continuum models. Acc. Chem. Res. 2000, 33 (12), 889–97.

(26) Saltalamacchia, A.; Casalino, L.; Borisek, J.; Batista, V. S.; Rivalta, I.; Magistrato, A. Decrypting the Information Exchange Pathways across the Spliceosome Machinery. J. Am. Chem. Soc. 2020, 142 (18), 8403–8411.

(27) Casalino, L.; Palermo, G.; Spinello, A.; Rothlisberger, U.; Magistrato, A. All-atom simulations disentangle the functional dynamics underlying gene maturation in the intron lariat spliceosome. Proc. Natl. Acad. Sci. U. S. A. 2018, 115 (26), 6584–6589.

(28) Verkhivker, G. M.; Agajanian, S.; Oztas, D. Y.; Gupta, G. Comparative Perturbation-Based Modeling of the SARS-CoV-2 Spike Protein Binding with Host Receptor and Neutralizing Antibodies: Structurally Adaptable Allosteric Communication Hotspots Define Spike Sites Targeted by Global Circulating Mutations. BioRxiv 2021.

(29) Nelson, G.; Buzko, O.; Spilman, P.; Niazi, K.; Rabizadeh, S.; Soon-Shiong, P. Molecular dynamic simulation reveals E484K mutation enhances spike RBD-ACE2 affinity and the combination of E484K, K417N and N501Y mutations (501Y.V2 variant) induces conformational change greater than N501Y mutant alone, potentially resulting in an escape mutant. BioRxiv 2021.

(30) Ray, D.; Le, L.; Andricioaei, I. Distant Residues Modulate Conformational Opening in SARS-CoV-2 Spike Protein. BioRxiv 2021.

(31) Van Wart, A. T.; Durrant, J.; Votapka, L.; Amaro, R. E. Weighted Implementation of Suboptimal Paths (WISP): An Optimized Algorithm and Tool for Dynamical Network Analysis. J. Chem. Theory Comput. 2014, 10 (2), 511–517.

(32) Wu, N. C.; Yuan, M.; Liu, H.; Lee, C. D.; Zhu, X.; Bangaru, S.; Torres, J. L.; Caniels, T. G.; Brouwer, P. J. M.; van Gils, M. J.; Sanders, R. W.; Ward, A. B.; Wilson, I. A. An Alternative Binding Mode of IGHV3-53 Antibodies to the SARS-CoV-2 Receptor Binding Domain. Cell Rep. 2020, 33 (3), 108274.

(33) Barnes, C. O.; Jette, C. A.; Abernathy, M. E.; Dam, K. A.; Esswein, S. R.; Gristick, H. B.; Malyutin, A. G.; Sharaf, N. G.; Huey-Tubman, K. E.; Lee, Y. E.; Robbiani, D. F.; Nussenzweig, M. C.; West, A. P., Jr.; Bjorkman, P. J. SARS-CoV-2 neutralizing antibody structures inform therapeutic strategies. Nature 2020, 588 (7839), 682–687.

(34) Ju, B.; Zhang, Q.; Ge, J.; Wang, R.; Sun, J.; Ge, X.; Yu, J.; Shan, S.; Zhou, B.; Song, S.; Tang, X.; Yu, J.; Lan, J.; Yuan, J.; Wang, H.; Zhao, J.; Zhang, S.; Wang, Y.; Shi, X.; Liu, L.; Zhao, J.; Wang, X.; Zhang, Z.; Zhang, L. Human neutralizing antibodies elicited by SARS-CoV-2 infection. Nature 2020, 584 (7819), 115–119.

(35) Wu, Y.; Wang, F.; Shen, C.; Peng, W.; Li, D.; Zhao, C.; Li, Z.; Li, S.; Bi, Y.; Yang, Y.; Gong, Y.; Xiao, H.; Fan, Z.; Tan, S.; Wu, G.; Tan, W.; Lu, X.; Fan, C.; Wang, Q.; Liu, Y.; Zhang, C.; Qi, J.; Gao, G. F.; Gao, F.; Liu, L. A noncompeting pair of human neutralizing antibodies block COVID-19 virus binding to its receptor ACE2. Science 2020, 368 (6496), 1274–1278.

(36) Barnes, C. O.; West, A. P., Jr.; Huey-Tubman, K. E.; Hoffmann, M. A. G.; Sharaf, N. G.; Hoffman, P. R.; Koranda, N.; Gristick, H. B.; Gaebler, C.; Muecksch, F.; Lorenzi, J. C. C.; Finkin, S.; Hagglof, T.; Hurley, A.; Millard, K. G.; Weisblum, Y.; Schmidt, F.; Hatziioannou, T.; Bieniasz, P. D.; Caskey, M.; Robbiani, D. F.; Nussenzweig, M. C.; Bjorkman, P. J. Structures of Human Antibodies Bound to SARS-CoV-2 Spike Reveal Common Epitopes and Recurrent Features of Antibodies. Cell 2020, 182 (4), 828–842 e16.

(37) Liu, L.; Wang, P.; Nair, M. S.; Yu, J.; Rapp, M.; Wang, Q.; Luo, Y.; Chan, J. F.; Sahi, V.; Figueroa, A.; Guo, X. V.; Cerutti, G.; Bimela, J.; Gorman, J.; Zhou, T.; Chen, Z.; Yuen, K. Y.; Kwong, P. D.; Sodroski, J. G.; Yin, M. T.; Sheng, Z.; Huang, Y.; Shapiro, L.; Ho, D. D. Potent neutralizing antibodies against multiple epitopes on SARS-CoV-2 spike. Nature 2020, 584 (7821), 450–456.

