## Supporting Information for "Allosteric Cross-Talk Among SARS-CoV-2 Spike’s Receptor-Binding Domain Mutations Triggers an Effective Hijacking of Human Cell Receptor"

Angelo Spinello,[a] Andrea Saltalamacchia,[b] Jure Boríšek[c] and Alessandra Magistrato[a,*]

a CNR-IOM c/o SISSA, via Bonomea 265, 34136 Trieste, Italy
b International School for Advanced Studies SISSA, via Bonomea 265, 34136 Trieste, Italy
c National Institute of Chemistry, Hajdrihova ulica 19, 1000 Ljubljana, Slovenia

**Supplementary Methods** Pag. 2-4

**Figure S1 to S8** Pag. 5-12

**Table S1 to S5** Pag. 13-18

**References** Pag. 19-20

### Supplementary Methods

#### Model Building and Classical Molecular Dynamics

The model of the adduct between angiotensin-converting enzyme 2 (ACE2) and the receptor-binding domain (RBD) of SARS-CoV-2 (RBD/ACE2) was taken from our previous study[1] and was based on a crystal structure reported at the beginning of the SARS-CoV-2 pandemic (pdb ID 6M0J).[2] We have introduced in this model the RBD mutations N501Y, E484K and K417N, typical of the South African (SA) variant (hereafter referred as $^{SA}$RBD/ACE2). Next, we have built three distinct RBD/ACE2 adducts carrying singularly the N501Y ($^{N501Y}$RBD/ACE2 or $^{UK}$RBD/ACE2), E484K ($^{E484K}$RBD/ACE2) and K417N ($^{K417N}$RBD/ACE2) mutations and we compared each model with the wild type RDB/ACE2 adduct. By deleting the ACE2 protein from these adducts, five isolated RBD were also generated (RBD, $^{SA}$RBD, $^{N501Y}$RBD, $^{E484K}$RBD, $^{K417N}$RBD).

The topologies for all the aforementioned models were built using the Parm99SB-ILDN Amber force field (FF) for the protein,[3] using the leap module of AmberTools 18.[4] The charge of the adducts was neutralized with the addition of $Na^+$ ions using the parameters of Joung and Cheatham.[5] All models were embedded in a 25 x 25 x 12 Å layer of TIP3P water molecules[6] leading to a box size of 117 x 127 x 140 Å$^3$, for a total of about 190,000 atoms. Afterwards, the topologies were converted to the GROMACS format using the acpype software.[7]

Classical Molecular Dynamics (MD) simulations were performed using the GROMACS 2020.2 software.[8] An integration time step of 2 fs was used and all covalent bonds involving H atoms were constrained with the LINCS algorithm. Particle Mesh Ewald (PME) scheme was used to account for electrostatic interactions, using a real space cut-off of 10 Å.[9] MD simulations were performed in the isothermal-isobaric NPT ensemble, at a temperature of 300 K, using a velocity-rescaling thermostat[10] and a Parrinello-Rahman barostat.[11]

For all systems, a preliminary energy minimization was performed using a steepest descent algorithm. Next, we gradually heated the system to 300 K with an increase of 50 K every 2 ns for a total of 12 ns, keeping the entire system highly restrained except for the solvent and solute hydrogens. Then, we switched to the NPT ensemble, scaling the pressure to 1 bar and using two different barostats: (i) the Berendsen barostat was used for 20 ns with the same restraints on the atoms and (ii) the Parrinello-Rahman barostat for 30 additional ns, while leaving the side chains free of constraint. Subsequently, we gradually decreased the restraints in 20 ns. Finally, each RBD/ACE2 adducts and isolated RBD models were relaxed for 2.5 μs and 500 ns of classical MD, respectively, leading to a cumulative simulation time of 15 μs of all-atom MD. From these trajectories, the last 2000 and 300 ns of the RBD/ACE2 adducts and isolated RBD models were respectively used for subsequent analyses.

**Analysis**

Root mean square fluctuation (RMSF), root mean square displacement (RMSD), hydrogen (H)-bond and Θ angle α1-helix of ACE2 analyses were performed using the *cpptraj* module of AmberTools 18 and GROMACS 2020.2 tools.[4,8]

The binding free energies ($\Delta G_b$) and their components were calculated with the Molecular Mechanics/Generalized Born Surface Area (MM-GBSA) method by using MMPBSA.py program on 100 frames taken from the equilibrated part of the trajectories.[12] Moreover, a per-residue decomposition analysis has been done.

**Principal Component Analysis (PCA)**

Principal Component Analysis has been performed by using the *cpptraj* module of Amber 18 to extract the coupled motions of the wild type and mutated SARS-CoV-2's RBD in complex with ACE2. This analysis allows identifying the collective motions of biological macromolecules, providing information on the most relevant conformational changes that occur during MD simulations. Through the diagonalization of the atomic position covariance matrix, it is possible to obtain a full set of eigenvectors, or Principal Components (PCs), and eigenvalues, reflecting the main motions of the protein and the associated amount of fluctuation, respectively.

**Correlation Coefficients (CCs)**

The cross-correlation matrices (or normalized covariance matrices), based on the Pearson's coefficient have been calculated on the equilibrated trajectories by using the *cpptraj* module of AmberTools 18. The analysis of the cross-correlation matrices allows determining qualitatively the coupling motions between two residues during the trajectory. These correlations have been detected by calculating the mass-weighted covariance matrix of Cα atoms' atomic fluctuations, after removing the global rotational and translational motions by fitting the structure to the initial frame of the equilibrated trajectory.

The element *ij* in the covariance matrix is the covariance between the atoms *i* and *j*:

$$C_{ij} = \langle (\vec{r}_i - \langle \vec{r}_i \rangle)(\vec{r}_j - \langle \vec{r}_j \rangle) \rangle$$

where $\vec{r}_i$ and $\vec{r}_j$ are the position vectors of atoms *i* and *j*, and the brackets denote an average over the sampled time period. A positive sign of this product indicates that the two atoms move in a correlated lockstep manner, while a negative value indicates an anticorrelated motion between the two atoms. If the product is zero, the displacements of the atoms are independent of each other.

The normalized covariance matrix by using the Pearson's coefficient gives the cross-correlation matrix (CCM) where each element *ij* corresponds to the Pearson's correlation coefficient (CC*ij*), between residue *i* and *j*. CC*ij* ranges from a value of -1, indicative of a totally anti-correlated motion between the two residues, to a value of +1, representative of a correlated motion.

**Dynamical Network Theory Analysis**

Allosteric paths were investigated using the Weighted Implementation of Suboptimal Paths (WISP) software,[13] employing the dynamical NetWork Analysis (NWA) to find cooperation between protein residues.

NWA enables finding the optimal and suboptimal communications pathways, which also greatly contribute to allosteric cross-talk and provide information about the quality and robustness of the signaling route. In dynamical network theory the protein is represented as a correlation-based weighted network, in which the nodes (center of mass of each residue), are connected by edges weighted by -log CC, which reflects the amount of correlations between each pair of residues. Hence, small/large weights indicate highly/poorly correlated and anticorrelated motions. After computing cross-correlations between residues along a MD trajectory NWA finds the optimal (i.e. shortest distance) through the weighted edges, and the slightly longer suboptimal communicating paths. A source and a sink residue, defining the beginning and the end of each path, need to be defined. The resulting paths have lengths defined in the network space which are inversely proportional to the amount of correlation existing among their constituent nodes.

For each trajectory, 5000 frames were taken (every 400 ps) from the equilibrated part of the MD simulation. These frames are then used by the WISP algorithm[13] to reconstruct the optimal path of correlated motions, along with 1000 suboptimal paths, which also contribute to the allosteric signaling across the RBD/ACE2 interface. For all the investigated adduct, the source and sink were represented by the amino acids in the position 484 and 501 of the RBD, respectively.

### Supplementary Figures

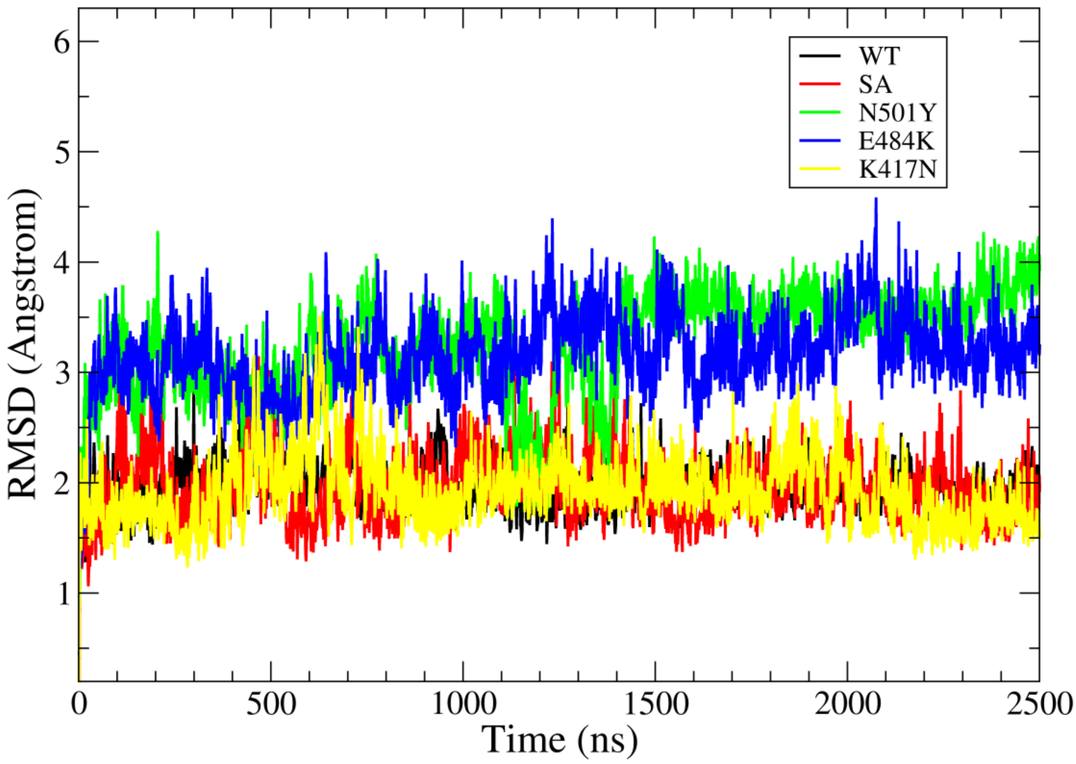


**Figure S1.** Root mean square deviation (RMSD, Å) vs simulation time (ns) calculated for the different receptor-binding domain (RBD) SARS-CoV-2 variants in complex with the angiotensin-converting enzyme 2 (ACE2) simulated in this work. The RBD/ACE2 models containing the wild type (WT), and South African (SA, i.e. N501Y and E484K and K417N), N501Y, E484K, and K417N RBD mutations are shown in black, red, green, blue and yellow lines, respectively.

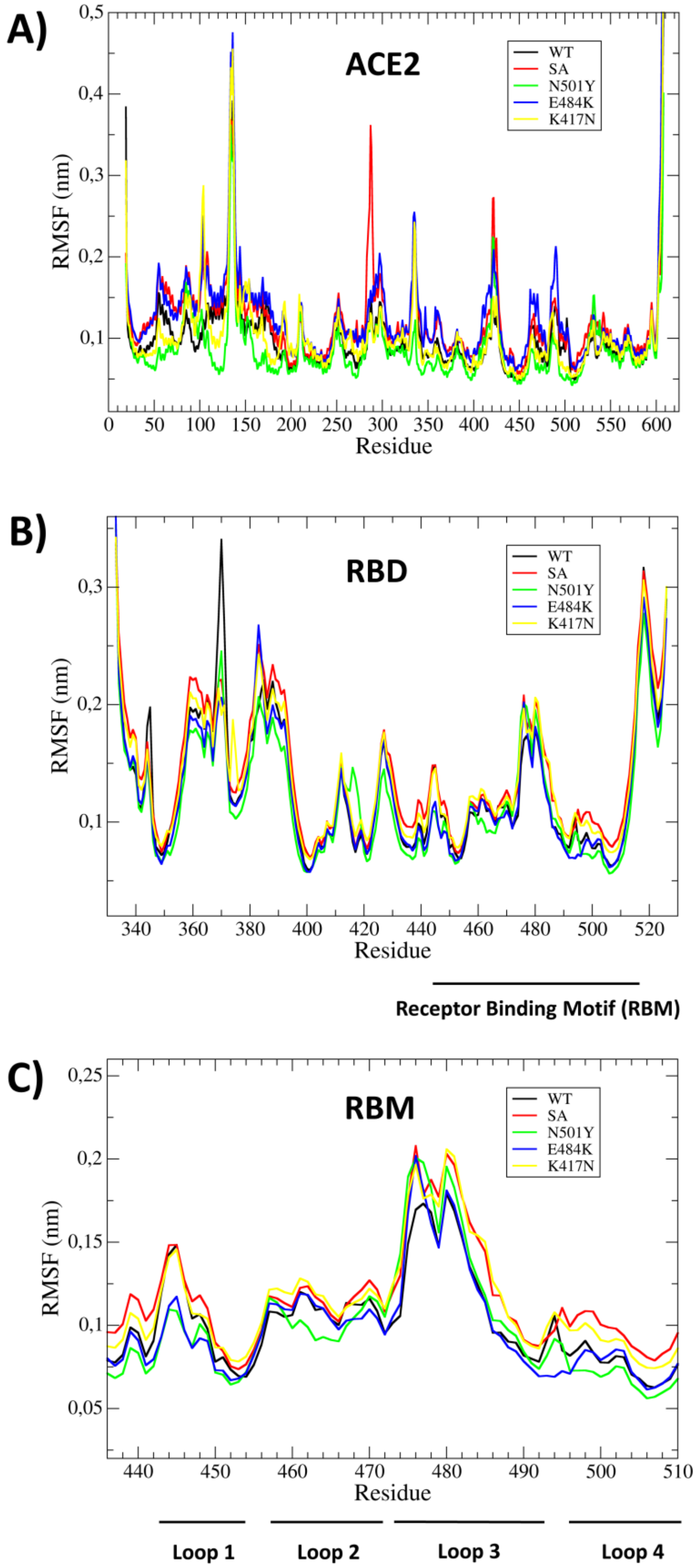


**Figure S2.** Root mean square fluctuation (RMSF) per residue (Å) calculated for A) the angiotensin-converting enzyme 2 (ACE2) receptor, B) the Spike's receptor-binding domain (RBD) and C) the receptor-binding motif (RBM). The RBD/ACE2 models containing the wild type (WT), South African (SA i.e. N501Y and E484K and K417N), N501Y, E484K, and K417N RBD mutations, are shown in black, red, green, blue and yellow lines, respectively.

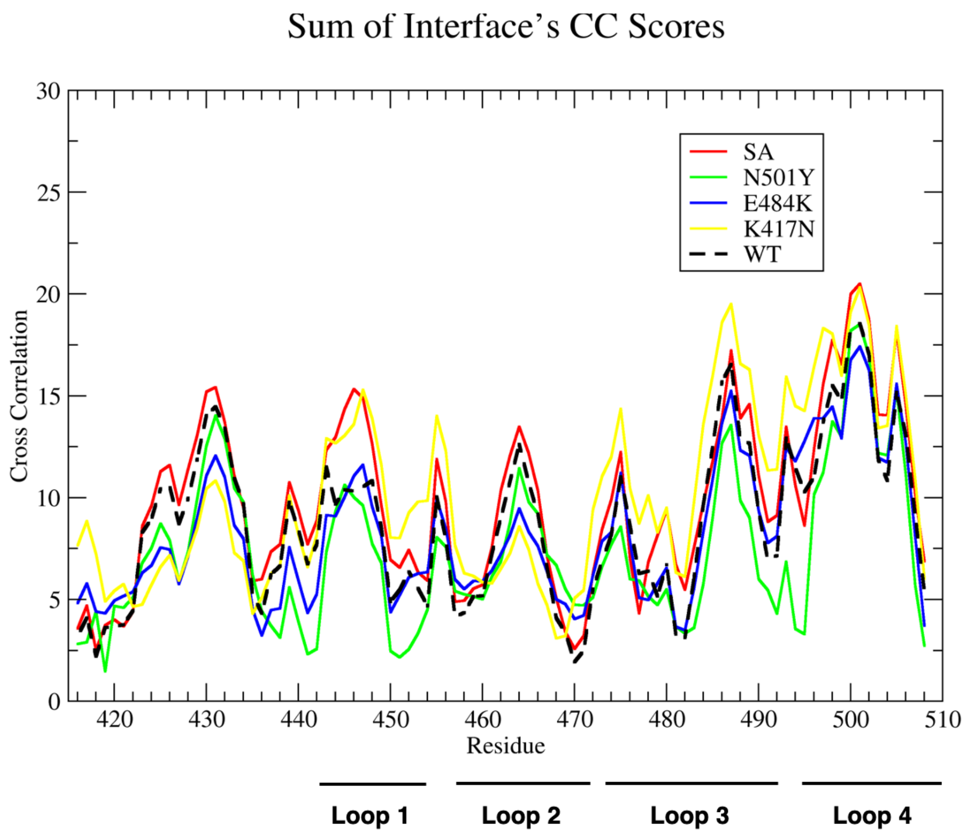


**Figure S3.** Sum of the per-residue cross-correlation coefficients for the receptor-binding motif (RBM)/angiotensin-converting enzyme 2 (ACE2) interface in the distinct Spike's receptor-binding domain (RBD)/ACE2 models investigated. The correlation of RBM residues of RBD/ACE2 models containing the wild type (WT), South African (SA i.e. N501Y and E484K and K417N), N501Y, E484K, and K417N RBD mutations, are shown in black, red, green, blue and yellow lines.

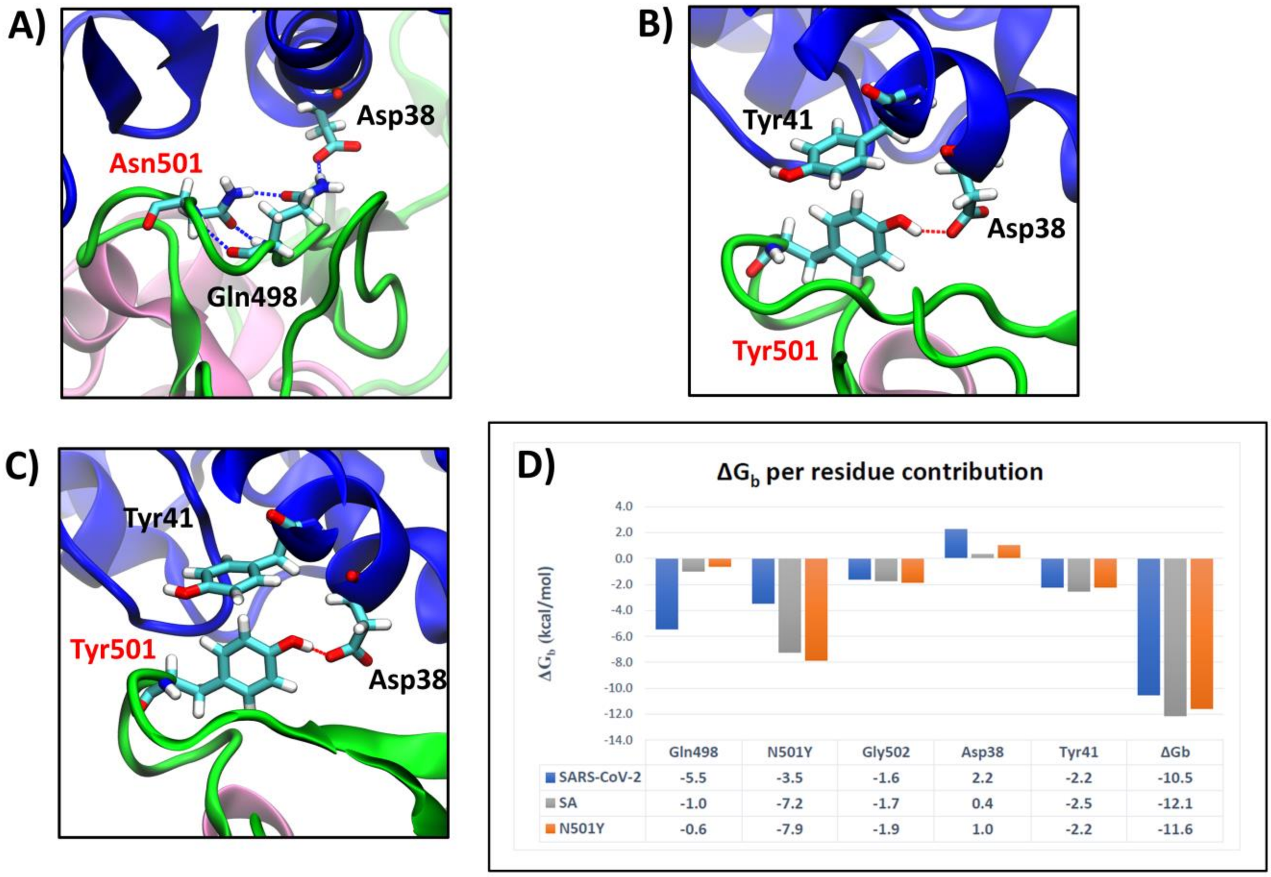


**Figure S4.** The N501Y mutations site in A) wild type receptor-binding domain (RBD)/angiotensin-converting enzyme 2 (ACE2) complex; B) in the $^{SA}$RBD/ACE2 and C) $^{N501Y}$RBD/ACE2 containing the South African set of RBD substitutions (N501Y, K417K, and E484K) and only the N501Y mutation, respectively. D) Per-residue binding free energy ($\Delta G_b$) calculated with the Molecular Mechanics/Generalized Born Surface Area (MM-GBSA) method.[12] ACE2, the RBM and RBD are shown as blue, green and pink new-cartoons, respectively, with interacting residues shown in licorice and colored by atom name.

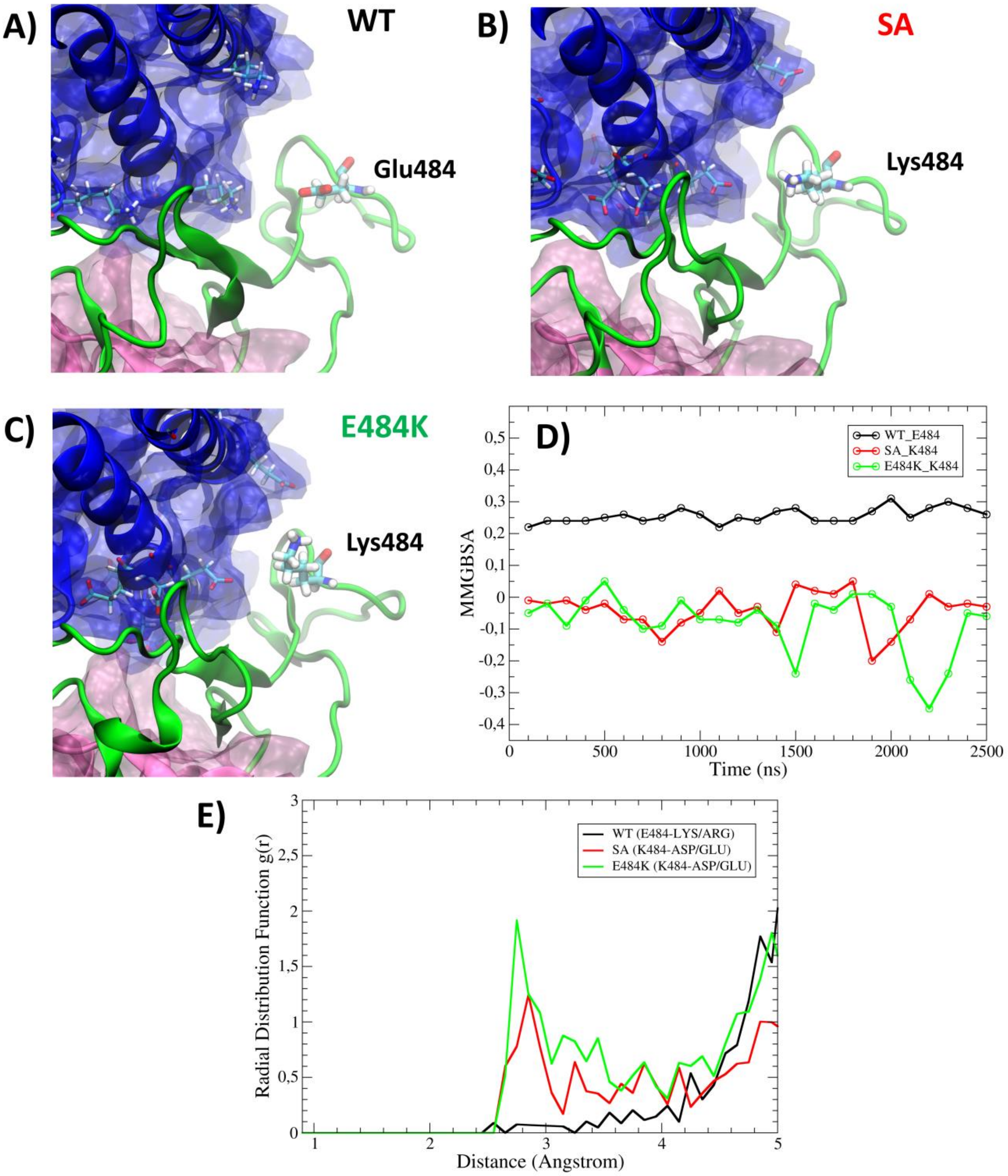


**Figure S5.** Representative structures as obtained from molecular dynamics (MD) trajectories of the receptor-binding domain (RBD)/angiotensin-converting enzyme 2 (ACE2) containing: A) wild type (WT) RBD/ACE2, B) the South African ($^{SA}$RBD/ACE2) set of RBD mutations (N501Y, E484K and K417K) and C) only the E484K mutation ($^{E484K}$RBD/ACE2) depicting the position of residue K484 and nearby negatively charged residues. ACE2, the RBM and RBD are shown as blue, green and pink new cartoons, respectively. D) Per-residue decomposition of the binding free energy ($\Delta G_b$) calculated with the Molecular Mechanics/Generalized Born Surface Area (MM-GBSA) method[12] for residue 484 in the WT RBD/ACE2 (black), $^{SA}$RBD/ACE2 (red) and $^{E484K}$RBD/ACE2 (green) plotted versus MD simulation time (ns). E) The radial distribution function, g(r), of residue 484 with respect to residues having an opposite charge.

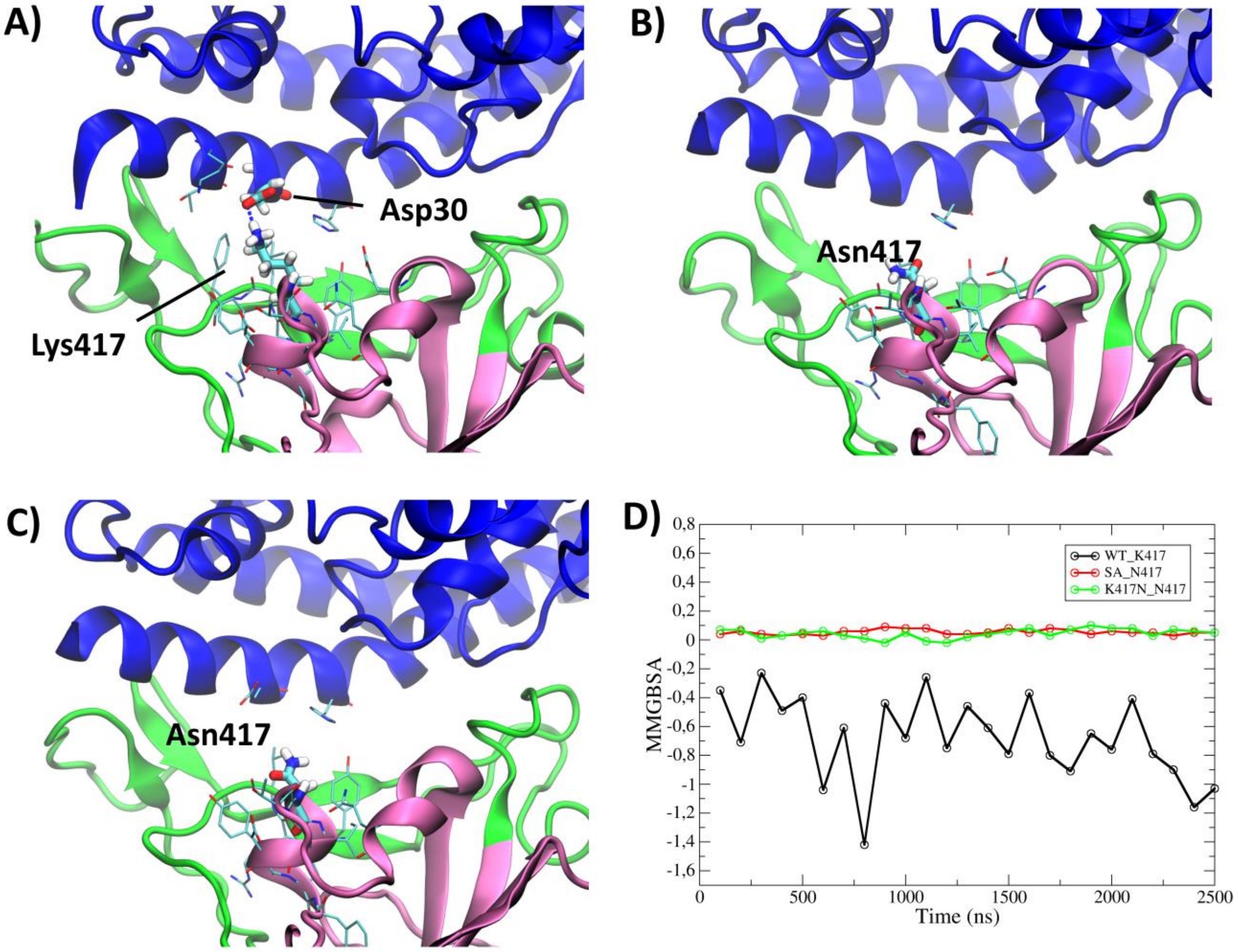


**Figure S6.** Representative structures showing the position of residue 417 as extracted from our molecular dynamics (MD) trajectory on A) wild type Spike's receptor-binding domain (RBD)/angiotensin-converting enzyme 2 (ACE2) adduct, B) $^{SA}$RDB/ACE2 and C) $^{K417N}$RBD/ACE2 adducts containing the South African (SA) set of RBD mutations (N501Y, E484K, and K417N) and only the K417 mutation, respectively. ACE2, the RBM and RBD are shown as blue, green and pink new-cartoons, respectively. D) Per-residue decomposition of the binding free energy ($\Delta G_b$, kcal/mol) for the residue 417 versus the MD simulation time (ns) calculated with the Molecular Mechanics/Generalized Born Surface Area (MM-GBSA) method[12] for the RBD/ACE2 (black), $^{SA}$RBD/ACE2 (red) and $^{K417N}$RBD/ACE2 (green) models.

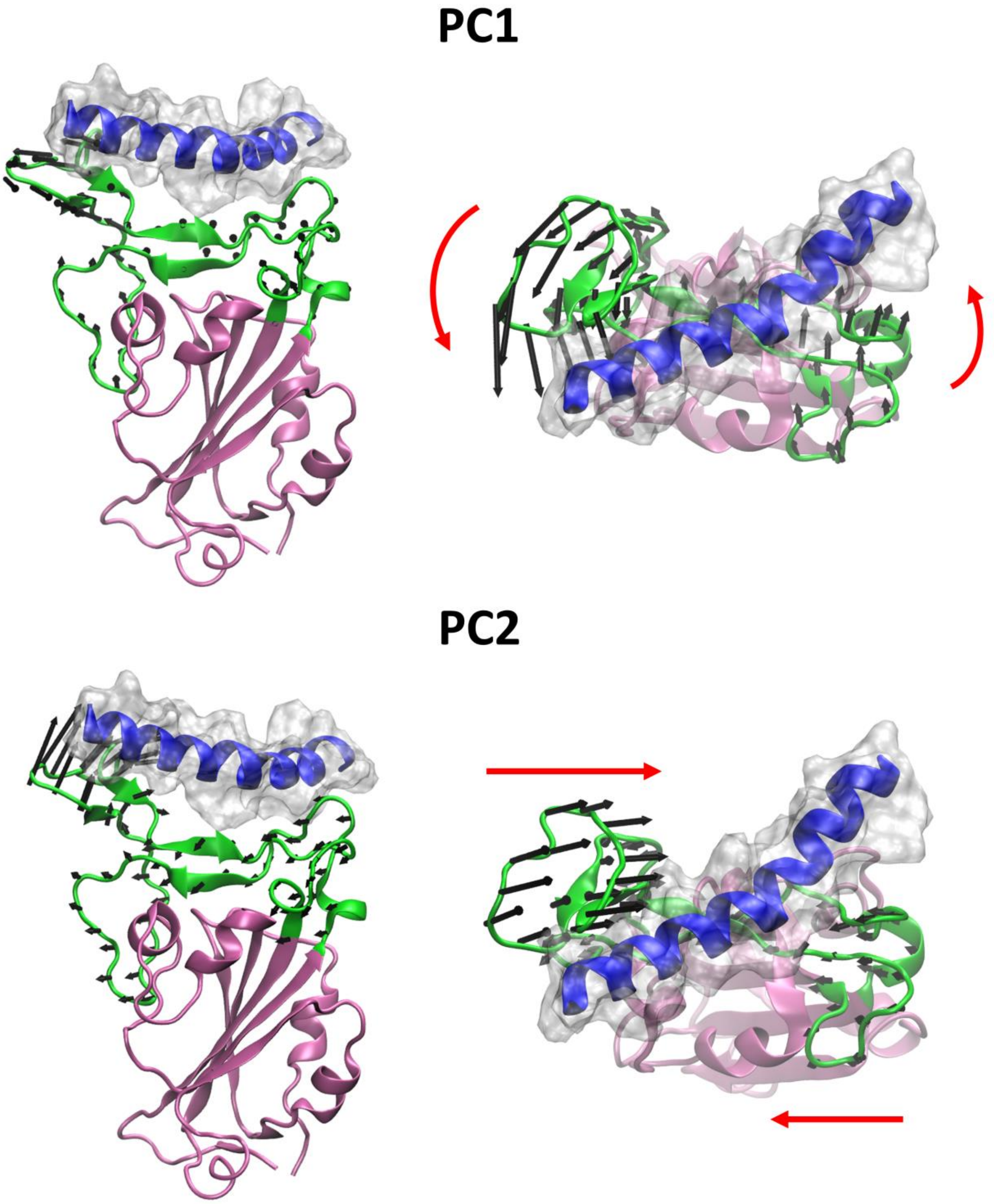


**Figure S7.** Front and top views of the two lowest-frequency principal component (PC1 and PC2), as extracted from the molecular dynamics (MD) simulations, of the isolated wild type receptor-binding domain (RBD). The receptor-binding motif (RBM) and RBD are shown as green and pink new-cartoons, respectively. Although being absent in the model the α1-helix of the angiotensin-converting enzyme 2 (ACE2) is highlighted as blue ribbons and a white transparent surface to better highlight how both PCs motions contribute to grasp the ACE2's α1-helix.

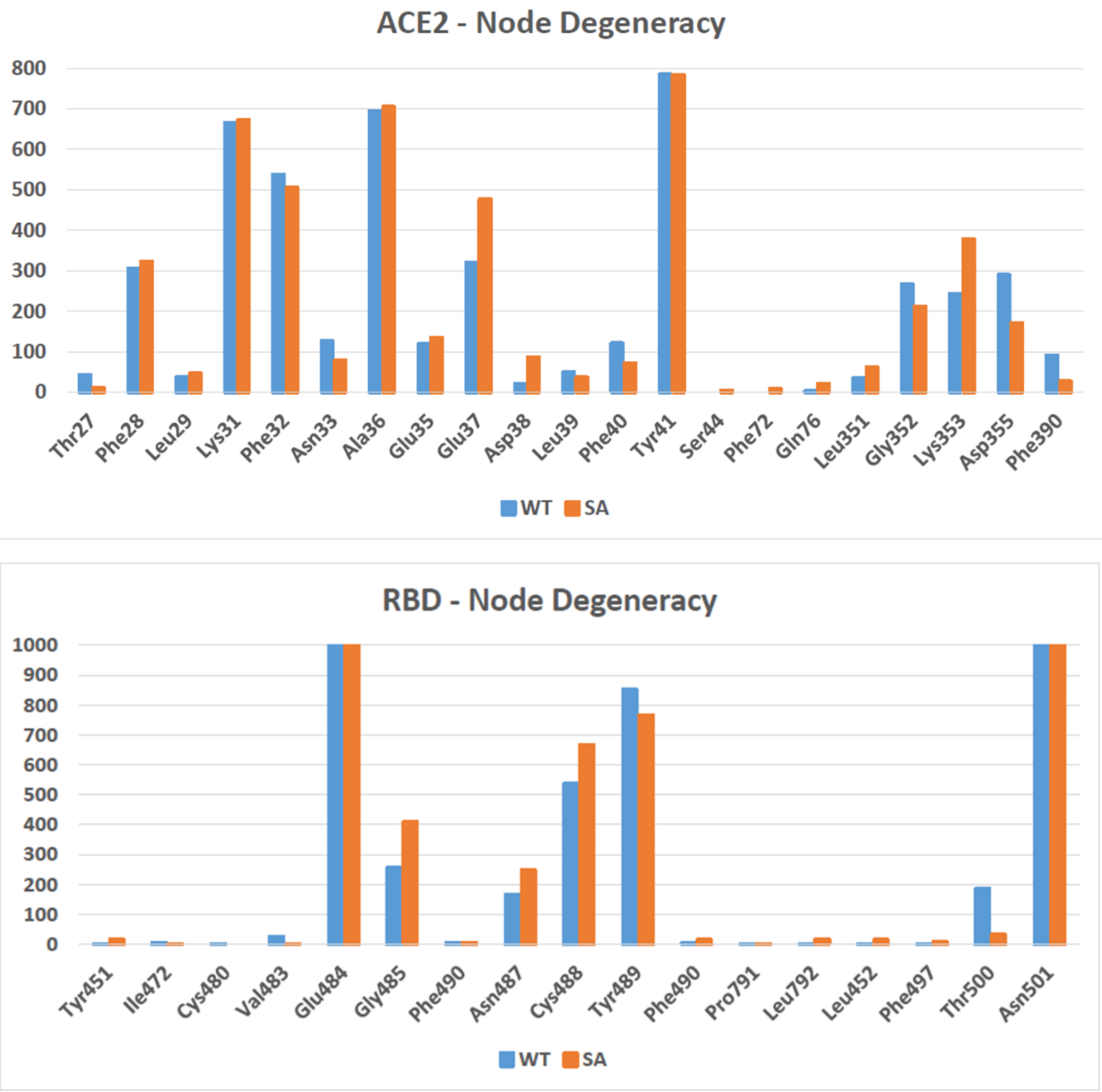


**Figure S8.** Node degeneracy in communication pathways. The total number of times a residue is crossed in the 1000 suboptimal pathways generated by the WISP program[13] is shown for wild type receptor-binding domain (RBD)/angiotensin-converting enzyme 2 (ACE2) (blue) and $^{SA}$RBD/ACE2 (orange) containing the South African (SA) set of mutations (i.e. N501Y, K417N, E484K). Source and sink are always residues 484 and 501, respectively.

### Supplementary Tables

**Table S1.** Persistency (%) and length (Å) of the hydrogen bonds forming at the receptor-binding domain (RBD)/angiotensin-converting enzyme 2 (ACE2) interface, as observed during the molecular dynamics simulations of all the investigated RBD/ACE2 adducts. The standard deviation is also reported. RBD/ACE2, $^{SA}$RBD/ACE2, $^{N501Y}$RBD/ACE2, $^{E484K}$RBD/ACE2, $^{K417N}$RBD/ACE2, refer to adducts containing the wild type, South African (SA i.e. N501Y, E484K and K417N), N501Y, E484K, and K417N RBD mutations. The mutated N501Y and K417N residues are highlighted in yellow.

| | | **RBD/ACE2** | | **$^{SA}$RBD/ACE2** | | **$^{N501Y}$RBD/ACE2** | | **$^{E484K}$RBD/ACE2** | | **$^{K417N}$RBD/ACE2** | |
|---|---|---|---|---|---|---|---|---|---|---|---|
| **SARS-CoV-2 residues** | **ACE2 residues** | **Persistence (%)** | **Average distance (Å)** | **Persistence (%)** | **Average distance (Å)** | **Persistence (%)** | **Average distance (Å)** | **Persistence (%)** | **Average distance (Å)** | **Persistence (%)** | **Average distance (Å)** |
| Thr500 | Asp355 | 92.4 | 2.68 | 82.2 | 2.68 | 93.3 | 2.68 | 67.8 | 2.72 | 88.4 | 2.69 |
| Asn487 | Tyr83 | 82.1 | 2.75 | 75.9 | 2.75 | 81.6 | 2.74 | 80.5 | 2.75 | 76.9 | 2.75 |
| Gln493 | Glu35 | 76.9 | 2.83 | 75.5 | 2.83 | 79.2 | 2.83 | 78.3 | 2.82 | 75.7 | 2.83 |
| Gly502 | Lys353 | 76.5 | 2.86 | 67.2 | 2.79 | 72.9 | 2.87 | 71.7 | 2.87 | 76.3 | 2.86 |
| Gln498 | Lys353 | 70.9 | 2.79 | - | - | - | - | 76.2 | 2.82 | 79.9 | 2.81 |
| Gln498 | Asp38 | 67.2 | 2.83 | - | - | - | - | 62.3 | 2.82 | 72.5 | 2.82 |
| N501Y | Asp38 | (Asn) - | - | (Tyr) 66.0 | 2.64 | (Tyr) 74.8 | 2.64 | (Asn) - | - | (Asn) - | - |
| Gln493 | Lys31 | 63.2 | 2.79 | 64.1 | 2.79 | 60.8 | 2.79 | 60.7 | 2.79 | 65.9 | 2.79 |
| K417N | Asp30 | (Lys) 47.2 | 2.79 | (Asn) - | - | (Lys) 41.8 | 2.79 | (Lys) 51.0 | 2.78 | (Asn) - | - |
| Tyr449 | Asp38 | 45.0 | 2.68 | - | - | - | - | 44.2 | 2.71 | 66.2 | 2.69 |

**Table S2.** Binding free energy ($\Delta G_b$, kcal/mol) of all investigated receptor-binding domain (RBD)/angiotensin-converting enzyme 2 (ACE2) adducts along with van der Waals, electrostatic contribution, solvation free energy calculated using a Generalized Born (EGB) approach and the empirically calculated nonpolar contribution to the solvation free energy (ESurf), as obtained with the Molecular Mechanics/Generalized Born (MM-GBSA) method.[12] The standard deviation is also reported. RBD/ACE2, $^{SA}$RBD/ACE2, $^{N501Y}$RBD/ACE2, $^{E484K}$RBD/ACE2 and $^{K417N}$RBD/ACE2 refer to adducts containing the wild type, South African (SA i.e. N501Y, K417 and E484K), N501Y, E484K and K417N RBD mutations.

| Energy | Van der Waals | Electrostatic | EGB | ESurf | $\Delta G_{bind}$ MM-GBSA |
|---|---|---|---|---|---|
| **RBD/ACE2** | -93.4 ± 4.9 | -711.6 ± 38.1 | 754.7 ± 35.4 | -13.6 ± 0.5 | **-63.9 ± 6.1** |
| **$^{SA}$RBD/ACE2** | -97.7 ± 6.7 | -862.3 ± 35.3 | 919.4 ± 35.8 | -13.1 ± 0.7 | **-53.7 ± 6.0** |
| **$^{N501Y}$RBD/ACE2** | -97.8 ± 7.9 | -688.7 ± 44.3 | 748.8 ± 43.7 | -13.4 ± 0.9 | **-51.1 ± 7.8** |
| **$^{E484K}$RBD/ACE2** | -94.2 ± 5.5 | -1192.1 ± 58.2 | 1236.8 ± 55.9 | -13.2 ± 8.9 | **-62.8 ± 8.9** |
| **$^{K417N}$RBD/ACE2** | 95.3 ± 5.7 | -468.3 ± 32.2 | 515.8 ± 32.2 | 13.4 ± 0.5 | **61.2 ± 6.3** |

**Table S3.** Node degeneracy in signaling paths. The number of times a node is present in the calculated paths is reported for receptor-binding domain (RBD)/angiotensin-converting enzyme 2 (ACE2) adducts. RBD/ACE2 and $^{SA}$RBD/ACE2 referring to models containing the wild type and South African variant (SA, i.e. N501Y, K417N and E484K). Sink and source residues are highlighted in cyan.

| | **RBD/ACE2** | **$^{SA}$RBD/ACE2** |
|---|---|---|
| **ACE2 Nodes** | **Degeneracy** | **Degeneracy** |
| Phe28 | 306 | 323 |
| Lys31 | 665 | 672 |
| Phe32 | 538 | 505 |
| Asn33 | 128 | 79 |
| Ala36 | 695 | 705 |
| Glu35 | 119 | 135 |
| Glu37 | 321 | 476 |
| Asp38 | 22 | 87 |
| Phe40 | 121 | 72 |
| Tyr41 | 785 | 783 |
| Gly352 | 268 | 212 |
| Lys353 | 243 | 378 |
| Asp355 | 290 | 171 |
| Lys353 | 243 | 378 |
| Asp355 | 290 | 171 |
| Phe390 | 92 | 27 |
| **RBD Nodes** | **Degeneracy** | **Degeneracy** |
| Glu484 | 1000 | 1000 |
| Gly485 | 257 | 410 |
| Asn487 | 167 | 248 |
| Cys488 | 537 | 668 |
| Tyr489 | 852 | 766 |
| Thr500 | 186 | 32 |
| Asn501 | 1000 | 1000 |

**Table S4.** Persistency (%) of the intramolecular hydrogen (H)-bonds formed near the receptor-binding domain (RBD)/angiotensin-converting enzyme 2 (ACE2) adduct interface during the molecular dynamics simulations. RBD/ACE2, $^{SA}$RBD/ACE2 and $^{K417N}$RBD/ACE2 refer to adducts containing the wild type, South African (SA i.e. N501Y, K417 and E484K) and K417N RBD mutations. H-bonds differences of more than 10% are highlighted in cyan. Mutation sites are highlighted in yellow. Intra-RBD H-bonds are separated according to their location in Loop1 to Loop4 (RBM-Loop1/RBM-Loop4), H-bonds connecting Loop1 to Loop4 (RBM-Loop1-Loop4), the β-sheets inside the RBM (RBM - β-sheets) and finally the remaining part of the RBD.

| RBD residues | | RBD/ACE2 | $^{SA}$RBD/ACE2 | $^{K417N}$RBD/ACE2 |
|---|---|---|---|---|
| | | **Persistence (%)** | **Persistence (%)** | **Persistence (%)** |
| **RBM – Loop1** | | | | |
| Asp442@Oδ | Ser438@Hγ | 97.7 | 97.6 | 98.0 |
| Asp442@Oδ | Arg509@Hε | 95.4 | 94.6 | 95.6 |
| Asp442@Oδ | Arg509@HH | 90.8 | 90.5 | 91.3 |
| Asp442@Oδ | Tyr451@HH | 91.4 | 90.9 | 87.7 |
| Asp442@O | Asn448@Hδ | 82.7 | 86.6 | 78.9 |
| Ser438@O | Asp442@H | 80.3 | 78.1 | 80.0 |
| Ser438@O | Leu441@H | 60.1 | 57.1 | 58.0 |
| Asn439@O | Ser443@Hγ | 49.6 | 65.9 | 58.1 |
| Lys444@O | Gly447@H | 42.2 | 45.4 | 39.8 |
| **RBM – Loop2** | | | | |
| Ser469@O | Arg454@HH | 85.6 | 86.5 | 86.7 |
| Ser469@O | Arg454@HH | 59.3 | 54.9 | 59.3 |
| Asp420@O | Leu461@H | 85.3 | 85.8 | 85.3 |
| Asp467@OD | Ser469@Hγ | 85.1 | 90.9 | 92.1 |
| Asp467@O | Arg454@HH | 82.8 | 84.8 | 84.8 |
| Ser459@O | Arg457@HH | 79.8 | 79.5 | 79.5 |
| Trp353@O | Arg466@HH | 60.6 | 67.7 | 74.1 |
| Asp467@Oδ | Arg457@Hε | 71.6 | 71.1 | 81.8 |
| Lys424@O | Phe464@H | 43.2 | 38.9 | 35.7 |
| **RBM – Loop3** | | | | |
| Tyr489@O | Tyr473@H | 86.2 | 83.8 | 83.6 |
| Asn487@O | Ala475@H | 62.5 | 44.9 | 45.7 |
| Cys488@O | Gly485@H | 55.5 | 52.3 | 51.1 |
| Gly476@O | Gln474@H | 51.0 | - | - |
| Tyr473@O | Tyr489@H | 41.1 | 48.9 | 57.3 |
| E484K@Oε | Phe490@H | (Glu) 19.9 | (Lys) - | (Glu) 17.0 |
| **RBM – Loop4** | | | | |
| N501Y@O | Gln506@Hε | (Asn) 82.8 | (Tyr) 82.6 | (Asn) 80.4 |
| Gly502@O | Tyr505@H | 60.2 | 56.0 | 57.4 |
| Tyr508@O | Ile402@H | 60.0 | 59.1 | 56.1 |
| Gln506@O | Gly404@H | 51.5 | 61.6 | 55.4 |
| Gly504@O | Asp505@H | 38.9 | 51.5 | 51.6 |
| Gln498@Oε | N501Y@Hδ | (Asn) 46.5 | (Tyr) - | (Asn) 49.3 |
| Gln498@O | N501Y@H | (Asn) 19.4 | (Tyr) 8.6 | (Asn) 17.3 |
| N501Y@Oδ | Gln498@H | (Asn) 18.6 | (Tyr) 8.2 | (Asn) 18.6 |

| | | | | |
|---|---|---|---|---|
| **RBM – Loop1-Loop4** | | | | |
| Gln506@Oε | Asn437@Hδ | 80.1 | 76.9 | 76.6 |
| Pro507@O | Ser438@H | 57.0 | 55.3 | 58.8 |
| Pro499@O | Asn439@Hδ | 51.5 | 51.6 | 48.7 |
| **RBM - β-sheets** | | | | |
| Asn422@Oδ | Arg454@H | 93.8 | 94.6 | 94.8 |
| Tyr453@O | Gln493@H | 87.4 | 89.1 | 89.1 |
| Pro491@O | Leu455@H | 76.3 | 68.0 | 71.1 |
| Gln759@O | Tyr753@H | 63.0 | 60.8 | 60.0 |
| Arg454@O | Asn422@Hδ | 53.5 | 56.3 | 58.3 |
| **RBD** | | | | |
| K417N@O | Asn422@Hδ | (Lys) 62.9 | (Asn) 66.3 | (Asn) 66.8 |
| Ile418@O | Tyr423@H | 61.9 | 63.8 | 62.5 |
| K417N@O | Tyr421@H | (Lys) 24.1 | (Asn) 34.0 | (Asn) 27.9 |

**Table S5.** Classification of class 1 and class 2 antibodies along with their Protein Data Bank (PDB) entry code and the list of the most important intermolecular interactions established with SARS-CoV-2 Spike's receptor-binding domain (RBD). H-bond refers to hydrogen bond.

| **Class 1** | | | |
|---|---|---|---|
| **Antibody** | **Reference** | **Protein Data Bank (PDB) entry** | **Interactions with K417** |
| B38 | Wu, et al.[14] | 7BZ5 | H-bond with Tyr52 |
| CC12.3 | Yuan, et al.[15] | 6XC7 | Salt Bridge with Asp98;<br>H-bond with Tyr52 |
| C102 | Barnes, et al.[16] | 7K8M | H-bond with backbone Gly97 |
| C105 | Barnes, et al.[17] | 6XCN | Salt Bridge with Glu96 |
| COVA2-04 | Wu, et al.[18] | 7JMO | Salt Bridge with Glu97 |
| **Class 2** | | | |
| **Antibody** | **Reference** | **Protein Data Bank (PDB) entry** | **Interactions with E484** |
| C002 | Barnes, et al.[16] | 7K8S | Salt Bridge with Arg96;<br>H-bond with backbone Ser94 |
| C121 | Barnes, et al.[16] | 7K8Y | H-bond with Tyr33 |
| C144 | Barnes, et al.[16] | 7K90 | H-bonds with Tyr58 |
| COVA2-39 | Wu, et al.[18] | 7JMP | H-bonds with Thr56 |
| P2B-2F6 | Ju, et al.[19] | 7BWJ | H-bonds with Tyr34 |
| Ab2-4 | Liu, et al.[20] | 6XEY | H-bonds with Asn52, Ser54 |

#### References


(1) Spinello, A.; Saltalamacchia, A.; Magistrato, A. Is the Rigidity of SARS-CoV-2 Spike Receptor-Binding Motif the Hallmark for Its Enhanced Infectivity? Insights from All-Atom Simulations. *J. Phys. Chem. Lett.* **2020,** *11* (12), 4785-4790.

(2) Lan, J.; Ge, J.; Yu, J.; Shan, S.; Zhou, H.; Fan, S.; Zhang, Q.; Shi, X.; Wang, Q.; Zhang, L.; Wang, X. Structure of the SARS-CoV-2 spike receptor-binding domain bound to the ACE2 receptor. *Nature* **2020,** *581* (7807), 215-220.

(3) Lindorff-Larsen, K.; Piana, S.; Palmo, K.; Maragakis, P.; Klepeis, J. L.; Dror, R. O.; Shaw, D. E. Improved side-chain torsion potentials for the Amber ff99SB protein force field. *Proteins* **2010,** *78* (8), 1950-8.

(4) D.A. Case, T. A. D., T.E. Cheatham, III, C.L. Simmerling, J. Wang, R.E. Duke, R. Luo, R.C. Walker, W. Zhang, K.M. Merz, B. Roberts, S. Hayik, A. Roitberg, G. Seabra, J. Swails, A.W. Götz, I. Kolossváry, K.F. Wong, F. Paesani, J. Vanicek, R.M. Wolf, J. Liu, X. Wu, S.R. Brozell, T. Steinbrecher, H. Gohlke, Q. Cai, X. Ye, J. Wang, M.-J. Hsieh, G. Cui, D.R. Roe, D.H. Mathews, M.G. Seetin, R. Salomon-Ferrer, C. Sagui, V. Babin, T. Luchko, S. Gusarov, A. Kovalenko, and P.A. Kollman. *AMBER 18,* University of California: San Francisco, 2018.

(5) Joung, I. S.; Cheatham, T. E., 3rd. Determination of alkali and halide monovalent ion parameters for use in explicitly solvated biomolecular simulations. *J. Phys. Chem. B* **2008,** *112* (30), 9020-41.

(6) Jorgensen, W. L.; Chandrasekhar, J.; Madura, J. D. Comparison of simple potential functions for simulating liquid water. *J. Chem. Phys.* **1983,** *79* (2), 926.

(7) Sousa da Silva, A. W.; Vranken, W. F. ACPYPE - AnteChamber PYthon Parser interfacE. *BMC Res Notes* **2012,** *5*, 367.

(8) Van Der Spoel, D.; Lindahl, E.; Hess, B.; Groenhof, G.; Mark, A. E.; Berendsen, H. J. GROMACS: fast, flexible, and free. *J. Comput. Chem.* **2005,** *26* (16), 1701-18.

(9) Darden, T.; York, D.; Pedersen, L. Particle mesh Ewald: An N·log(N) method for Ewald sums in large systems. *J. Chem. Phys.* **1993,** *98* (12), 10089.

(10) Bussi, G.; Donadio, D.; Parrinello, M. Canonical sampling through velocity rescaling. *J. Chem. Phys.* **2007,** *126* (1), 014101.

(11) Parrinello, M.; Rahman, A. Polymorphic transitions in single crystals: A new molecular dynamics method. *J. Appl. Phys.* **1981,** *52*, 7182.

(12) Kollman, P. A.; Massova, I.; Reyes, C.; Kuhn, B.; Huo, S.; Chong, L.; Lee, M.; Lee, T.; Duan, Y.; Wang, W.; Donini, O.; Cieplak, P.; Srinivasan, J.; Case, D. A.; Cheatham, T. E., 3rd. Calculating structures and free energies of complex molecules: combining molecular mechanics and continuum models. *Acc. Chem. Res.* **2000,** *33* (12), 889-97.

(13) Van Wart, A. T.; Durrant, J.; Votapka, L.; Amaro, R. E. Weighted Implementation of Suboptimal Paths (WISP): An Optimized Algorithm and Tool for Dynamical Network Analysis. *J. Chem. Theory Comput.* **2014,** *10* (2), 511-517.

(14) Wu, Y.; Wang, F.; Shen, C.; Peng, W.; Li, D.; Zhao, C.; Li, Z.; Li, S.; Bi, Y.; Yang, Y.; Gong, Y.; Xiao, H.; Fan, Z.; Tan, S.; Wu, G.; Tan, W.; Lu, X.; Fan, C.; Wang, Q.; Liu, Y.; Zhang, C.; Qi, J.; Gao, G. F.; Gao, F.; Liu, L. A noncompeting pair of human neutralizing antibodies block COVID-19 virus binding to its receptor ACE2. *Science* **2020,** *368* (6496), 1274-1278.

(15) Yan, R.; Zhang, Y.; Li, Y.; Xia, L.; Guo, Y.; Zhou, Q. Structural basis for the recognition of SARS-CoV-2 by full-length human ACE2. *Science* **2020,** *367* (6485), 1444-1448.

(16) Barnes, C. O.; Jette, C. A.; Abernathy, M. E.; Dam, K. A.; Esswein, S. R.; Gristick, H. B.; Malyutin, A. G.; Sharaf, N. G.; Huey-Tubman, K. E.; Lee, Y. E.; Robbiani, D. F.; Nussenzweig, M. C.; West, A. P., Jr.; Bjorkman, P. J. SARS-CoV-2 neutralizing antibody structures inform therapeutic strategies. *Nature* **2020,** *588* (7839), 682-687.

(17) Barnes, C. O.; West, A. P., Jr.; Huey-Tubman, K. E.; Hoffmann, M. A. G.; Sharaf, N. G.; Hoffman, P. R.; Koranda, N.; Gristick, H. B.; Gaebler, C.; Muecksch, F.; Lorenzi, J. C. C.; Finkin, S.; Hagglof, T.; Hurley, A.; Millard, K. G.; Weisblum, Y.; Schmidt, F.; Hatziioannou, T.; Bieniasz, P. D.; Caskey, M.; Robbiani, D. F.; Nussenzweig, M. C.; Bjorkman, P. J. Structures of Human Antibodies Bound to SARS-CoV-2 Spike Reveal Common Epitopes and Recurrent Features of Antibodies. *Cell* **2020,** *182* (4), 828-842 e16.

(18) Wu, N. C.; Yuan, M.; Liu, H.; Lee, C. D.; Zhu, X.; Bangaru, S.; Torres, J. L.; Caniels, T. G.; Brouwer, P. J. M.; van Gils, M. J.; Sanders, R. W.; Ward, A. B.; Wilson, I. A. An Alternative Binding Mode of IGHV3-53 Antibodies to the SARS-CoV-2 Receptor Binding Domain. *Cell Rep.* **2020,** *33* (3), 108274.

(19) Ju, B.; Zhang, Q.; Ge, J.; Wang, R.; Sun, J.; Ge, X.; Yu, J.; Shan, S.; Zhou, B.; Song, S.; Tang, X.; Yu, J.; Lan, J.; Yuan, J.; Wang, H.; Zhao, J.; Zhang, S.; Wang, Y.; Shi, X.; Liu, L.; Zhao, J.; Wang, X.; Zhang, Z.; Zhang, L. Human neutralizing antibodies elicited by SARS-CoV-2 infection. *Nature* **2020,** *584* (7819), 115-119.

(20) Liu, L.; Wang, P.; Nair, M. S.; Yu, J.; Rapp, M.; Wang, Q.; Luo, Y.; Chan, J. F.; Sahi, V.; Figueroa, A.; Guo, X. V.; Cerutti, G.; Bimela, J.; Gorman, J.; Zhou, T.; Chen, Z.; Yuen, K. Y.; Kwong, P. D.; Sodroski, J. G.; Yin, M. T.; Sheng, Z.; Huang, Y.; Shapiro, L.; Ho, D. D. Potent neutralizing antibodies against multiple epitopes on SARS-CoV-2 spike. *Nature* **2020,** *584* (7821), 450-456.
